# MAESTRO affords ‘breadth and depth’ for mutation testing

**DOI:** 10.1101/2021.01.22.427323

**Authors:** Gregory Gydush, Erica Nguyen, Jin H. Bae, Justin Rhoades, Sarah C. Reed, Douglas Shea, Kan Xiong, Ruolin Liu, Timothy Blewett, Fangyan Yu, Ka Wai Leong, Atish D. Choudhury, Daniel G. Stover, Sara M. Tolaney, Ian E. Krop, J. Christopher Love, Heather A. Parsons, G. Mike Makrigiorgos, Todd R. Golub, Viktor A. Adalsteinsson

## Abstract

The ability to assay large numbers of low-abundance mutations is crucial in biomedicine. Yet, the technical hurdles of sequencing multiple mutations at extremely high depth and accuracy remain daunting. For sequencing low-level mutations, it’s either ‘depth or breadth’ but not both. Here, we report a simple and powerful approach to accurately track thousands of distinct mutations with minimal reads. Our technique called MAESTRO (<u>m</u>inor <u>a</u>llele <u>e</u>nriched <u>s</u>equencing <u>t</u>hrough <u>r</u>ecognition <u>o</u>ligonucleotides) employs massively-parallel mutation enrichment to empower duplex sequencing—one of the most accurate methods—to track up to 10,000 low-frequency mutations with up to 100-fold less sequencing. In example use cases, we show that MAESTRO could enable mutation validation from cancer genome sequencing studies. We also show that it could track thousands of mutations from a patient’s tumor in cell-free DNA, which may improve detection of minimal residual disease from liquid biopsies. In all, MAESTRO improves the breadth, depth, accuracy, and efficiency of mutation testing.

## Introduction

Mutations in DNA emerge from single cells^1^, define cell populations^2^, and establish genetic diversity^3^. Considering the vast genetic diversity of living organisms and the significance of mutations in disease biology^4^, there is a growing need to assay many distinct, low-abundance mutations in multiple areas of biomedicine spanning oncology^5^, obstetrics^6^, transplantation^7,8^, infectious disease^9^, genetics^10^, microbiomics^11^, forensics^12^, and beyond. Yet, the intrinsic tradeoff in breadth-versus-depth of DNA sequencing means that either few mutations can be assayed at high depth, or many mutations at low depth—not both. High depth (i.e. many reads per genomic locus) is required to accurately detect low-abundance mutations, but this severely limits breadth (i.e. number of distinct loci). This explains why, despite massive reductions in sequencing costs, it remains prohibitively expensive to test for large numbers of distinct, low-abundance mutations.

Duplex sequencing is one of the most accurate methods for mutation detection, with 1000-fold fewer errors than standard sequencing, but adds significant cost^13^. By requiring mutations to be present in replicate reads from both strands of each DNA duplex, many of the errors in sample preparation and sequencing can be overcome to enable reliable detection of low-abundance mutations. Yet, up to 100-fold more reads per locus are required—a challenge that is exacerbated when tracking many low-abundance mutations. Less stringent methods exist that require fewer reads, but compromising specificity to save cost would be deeply problematic for applications that impact patient care. While methods to enrich rare mutations have been developed^14–21^, none have employed high-accuracy sequencing, nor tracked many rare mutations.

Liquid biopsy represents an application for which accurate, low-cost tracking of many distinct mutations could empower clinical decisions^22^. For instance, applying liquid biopsies to detect minimal residual disease (MRD) after cancer treatment^5,23–26^ has the potential to inform whether surgery is needed after neoadjuvant therapy^27–29^, whether adjuvant therapy is needed after surgery^30^, and ultimately, whether it is safe to stop treatment^31^. It could also enable treatment response to be monitored over several log-fold-changes in cancer burden, which has been critical in hematologic malignancies^32^, but is not yet feasible for most patients due to limited sensitivity.

One promising way to improve sensitivity of liquid biopsies is to track many patient-specific tumor mutations in cell-free DNA (cfDNA), recognizing that not all mutations may be present in an individual blood tube when tumor DNA in the bloodstream is sparse (i.e. less than a genome equivalent of tumor DNA per tube)^27,28,33–35^. Yet, this has been challenging, because to rely upon any subset for MRD detection requires extremely accurate sequencing of many rare mutations. We reason that methods to deplete the normal (i.e. non-tumor derived) cfDNA could enable accurate, low-cost tracking of thousands of mutations in a patient’s tumor genome and improve MRD detection.

Here, we describe MAESTRO (<u>m</u>inor <u>a</u>llele <u>e</u>nriched <u>s</u>equencing <u>t</u>hrough <u>r</u>ecognition <u>o</u>ligonucleotides), a technique which combines massively-parallel mutation enrichment with duplex sequencing to enable accurate, low-cost mutation testing. In contrast to conventional hybrid-capture duplex sequencing^34,36,37^ (herein referred to as ‘Conventional’), which uses long probes to capture mutant and wild type with similar efficiency, MAESTRO uses short probes to enrich for patient-specific mutant alleles and uncovers the same mutant duplexes using up to 100-fold fewer reads. We first establish the performance of MAESTRO in dilution series. Then, we provide two proof-of-principle applications. In the first, we show that MAESTRO could enable verification of low-abundance mutations discovered from cancer whole-exome sequencing. In the second, we show that MAESTRO could enable thousands of mutations from a patient’s tumor to be assayed in cfDNA, which may improve the detection of MRD.

## Results

### MAESTRO uncovers the same mutant duplexes with ~100-fold less sequencing

We have established an accurate and efficient technique to track large numbers of low-abundance mutations in clinical specimens (**Fig 1A**). Our technique, called MAESTRO, utilizes allele-specific hybridization with short probes, leveraging thermodynamic differences in heteroduplex versus homoduplex DNA (**Supplementary Fig. 1**), to enrich barcoded library molecules bearing up to 10,000 prespecified mutations. Minimal sequencing is applied, and mutations are detected on both sense strands of each DNA duplex (**Fig 1B**). MAESTRO also employs a tunable noise filter which excludes error-prone loci (**Methods**).

**Figure 1:**
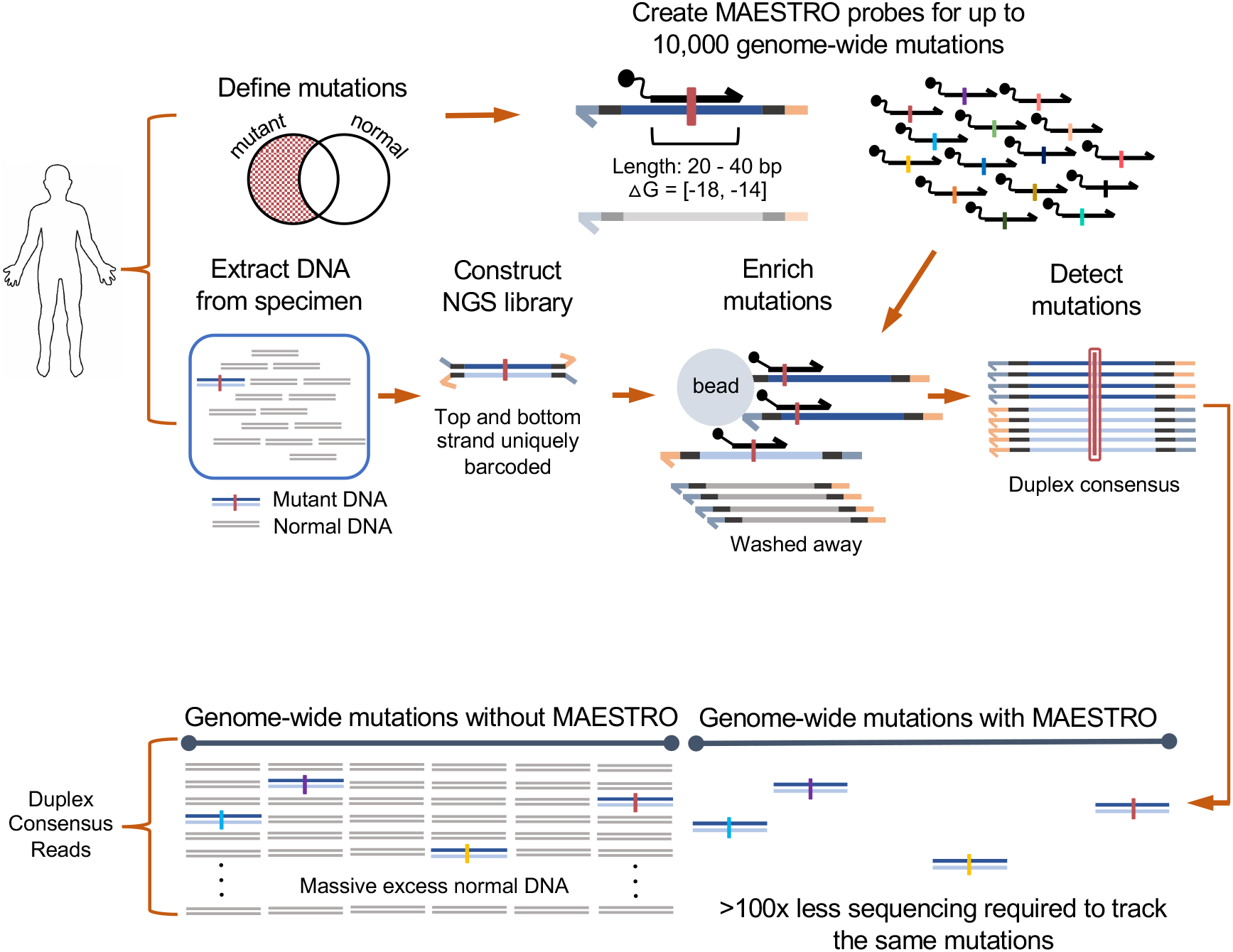
MAESTRO enables accurate, low-cost mutation tracking in clinical specimens. (A) Up to 10,000 MAESTRO probes are designed with stringent length and Δ*G* for single-nucleotide discrimination of predefined mutations (Supplementary Fig. 1, Methods). DNA libraries containing uniquely barcoded top and bottom strands are subject to hybrid capture using allele-specific MAESTRO probes. Only molecules containing tracked mutations are captured and sequenced with duplex consensus for error suppression. (B) Using MAESTRO, the same mutations are discovered using up to 100x less sequencing because uninformative regions are depleted.

We first sought to maximize fold-enrichment while minimizing loss of mutations. We created a 1/1k dilution of sheared genomic DNA from two human cell lines, identified exclusive single nucleotide polymorphisms (SNPs) as proxies for clonal mutations, and generated duplex sequencing libraries that were split for hybrid capture. Using qPCR, we confirmed that adapter ligation efficiencies are consistent with prior reports (**Supplementary Table 1**), and that MAESTRO capture efficiency is only slightly lower than conventional capture (35% vs. 57% respectively, **Supplementary Table 2**). After sequencing, we compared raw variant allele fraction (raw VAF) and recall of mutant duplexes (**Supplementary Table 3, Supplementary Fig. 3B)** using MAESTRO versus conventional hybrid capture (120 bp probes, 65°C annealing). By adjusting probe length and hybridization parameters, we established conditions (Δ*G* −18 to −14 kcal/mol, T=50°C, **Supplementary Fig. 1**, **Supplementary Fig. 2, Supplementary Fig. 3A**) that yielded strong fold-enrichment of mutant vs. wild type alleles (median 948.3-fold, range 8.1 to 3.4E4) while uncovering the majority of mutant duplexes (**Fig. 2A,B**, **Supplementary Table 3**). Indeed, the median raw VAF with MAESTRO was 0.97 (range 5.03E-3 to 1), in contrast to 6.98E-4 (range 3.00E-5 to 3.87E-3) with Conventional. The fraction of recoverable mutations (or, enrichment ‘success rate’) was 72.5%. Interestingly, we did not observe equal and opposite magnitude raw VAF changes when swapping strands of C and G reference base probes (**Supplementary Fig. 3C**). We believe this may be due to differences in probe characteristics (i.e. delta G, length) for each base category but further investigation is needed. MAESTRO cannot uncover more mutations than physically present in a sample; yet, by detecting each with up to 100x fewer reads, it can recover more total unique mutations, particularly when it would not otherwise be possible (e.g. due to cost) to sequence a sample to saturation.

**Figure 2:**
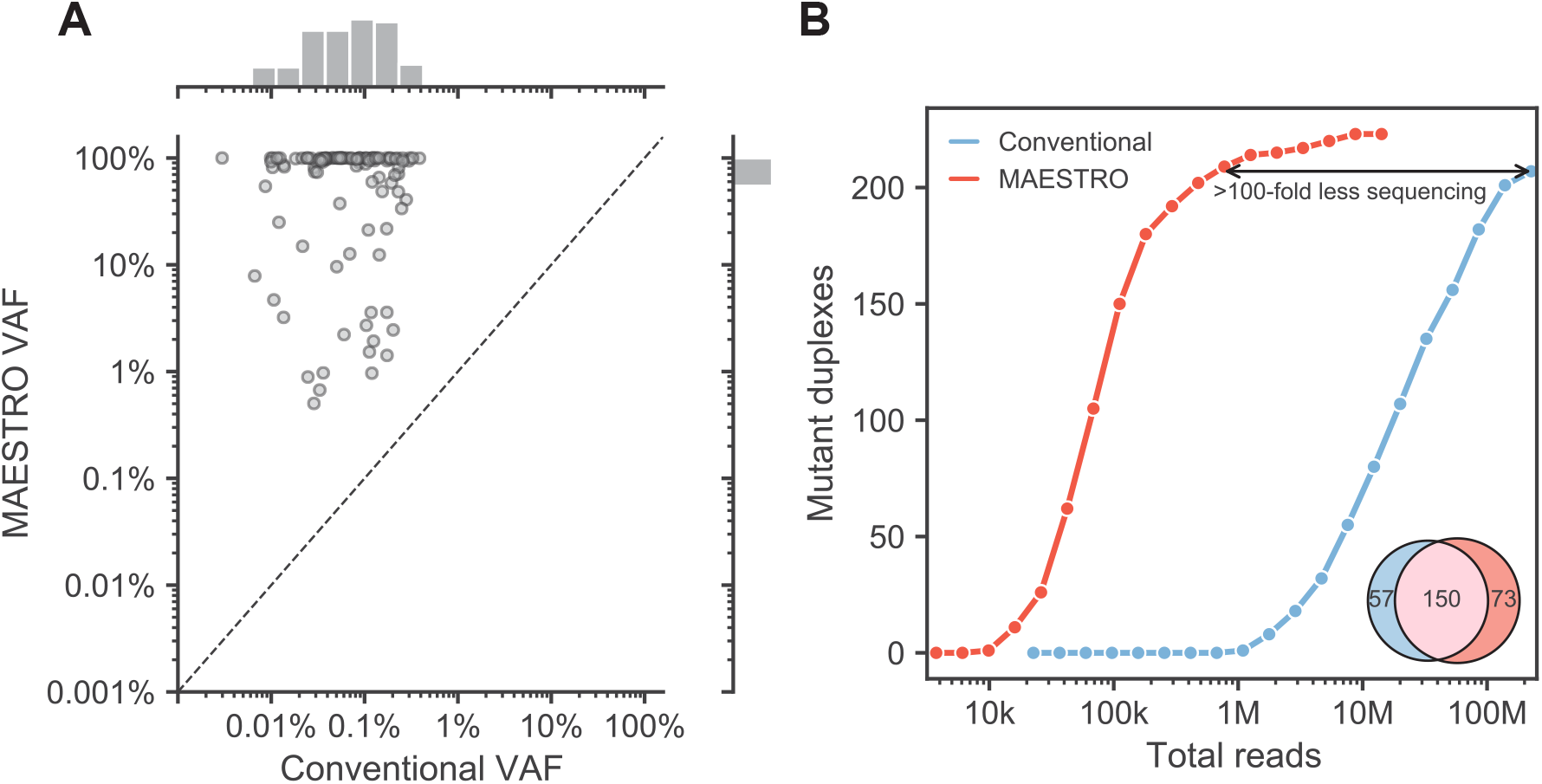
MAESTRO uncovers most mutant duplexes using significantly fewer reads. (A) Comparison of variant allele frequency with conventional duplex sequencing to MAESTRO with 438 probe panel at 1/1k tumor fraction. (B) Downsampling of conventional duplex sequencing and MAESTRO. As an inset, mutant duplex overlap is shown; of the 57 mutant duplexes exclusive to Conventional, 42 were detected by MAESTRO but excluded by the noise filter. The initial sample was barcoded with UMIs (unique molecular indices) which allowed for tracking individual duplex molecules through different experimental conditions.

We next tuned the MAESTRO noise filter. This filter was designed to protect against the possibility that errors could arise independently on both strands of library molecules and, given enrichment bias, ‘collide’ to form a duplex (**Supplementary Fig. 4A,B**). It works based on the assumptions that (i) errors should be impartial to read family, and (ii) error-prone loci should therefore exhibit a disproportionate number of double- (DSC) to single- (SSC) strand consensus read families bearing mutations (**Supplementary Fig. 4B**). Indeed, we found that sites with DSC/SSC ratios below 0.15 had poor reproducibility in replicate captures of a non-mutant library (our negative control) (**Supplementary Fig. 4C**). We also found that our filter protected against errors introduced by excessive PCR (**Supplementary Fig. 4D**), and further confirmed that MAESTRO probes—which contain the mutant base—do not create false mutant duplexes (**Supplementary Fig. 5**). Filtering by DSC/SSC ratio was found to be robust to changes in sequencing depth with similar concordance observed at 10% of the original sequencing depth (**Supplementary Fig. 6**).

Considering the profound enrichment, we then asked how many fewer reads would be required to detect the same mutant duplexes as Conventional. We found that MAESTRO could uncover the majority (n=150/207) using ~100-fold less sequencing (**Fig. 2B**), while providing comparable specificity (**Supplementary Fig. 7C**). Interestingly, of the 57 mutant duplexes exclusive to Conventional, 42 were detected by MAESTRO but excluded by the noise filter. Our results suggest that MAESTRO can uncover the majority of mutant duplexes using significantly less sequencing.

### MAESTRO enables mutation verification from tumor sequencing

Expansive methods such as whole-exome and whole-genome sequencing stand to unravel the genetic basis of human diseases. However, it remains challenging to resolve low-level mutations (e.g. < 10% VAF) given insufficient depth to read each DNA molecule enough times to suppress errors. Currently, mutations discovered in sequencing studies may be orthogonally validated via technologies such as digital droplet PCR or multiplex amplicon sequencing. However, these are not highly scalable approaches and are usually restricted to a handful of mutations suspected of having potential clinical significance. We reasoned that MAESTRO could enable rapid, low-cost verification of large numbers of mutations discovered from whole-exome and -genome sequencing. The net result would be that lower abundance mutations could be reliably discovered and verified from comprehensive sequencing studies.

To explore this, we performed whole-exome sequencing of tumor biopsies (of varied tumor purity; median 63%, range 26 - 100%) and matched normal DNA from 16 patients. We identified a median of 40 mutations per patient (median 40, range 13-130) and created both a MAESTRO and Conventional panel comprising all mutations for which we could design probes. Requiring the true mutations to be detected on both strands of each duplex, we found similar fractions of validated mutations between MAESTRO and Conventional, with slightly lower fractions for MAESTRO likely due to probe dropout (**Fig. 3A**). Yet, the fraction of validated mutations was much higher for those which had been identified at >0.10 VAF from tumor whole-exome sequencing (median 0.75, range 0.21-0.90 for MAESTRO; median 0.98, range 0.40-1.0 for Conventional), in comparison to those which had been identified at <0.10 VAF (median 0.29, range 0.07-0.82 for MAESTRO; median 0.35, range 0.04-1.0 for Conventional, **Fig. 3A**). Indeed, the mutations which were found to be “not validated” tended to have the lowest VAFs from tumor whole-exome sequencing (median 0.04, range 0.01-0.83, **Fig. 3B**). Expectedly, we observed higher fractions of MAESTRO-validated mutations in fresh-frozen (median 0.65, range 0.62-0.77) as compared to formalin-fixed (median 0.58, range 0.10-0.76) tumor biopsies. Our results suggest that MAESTRO could be an invaluable tool for validation in mutation discovery efforts.

**Figure 3:**
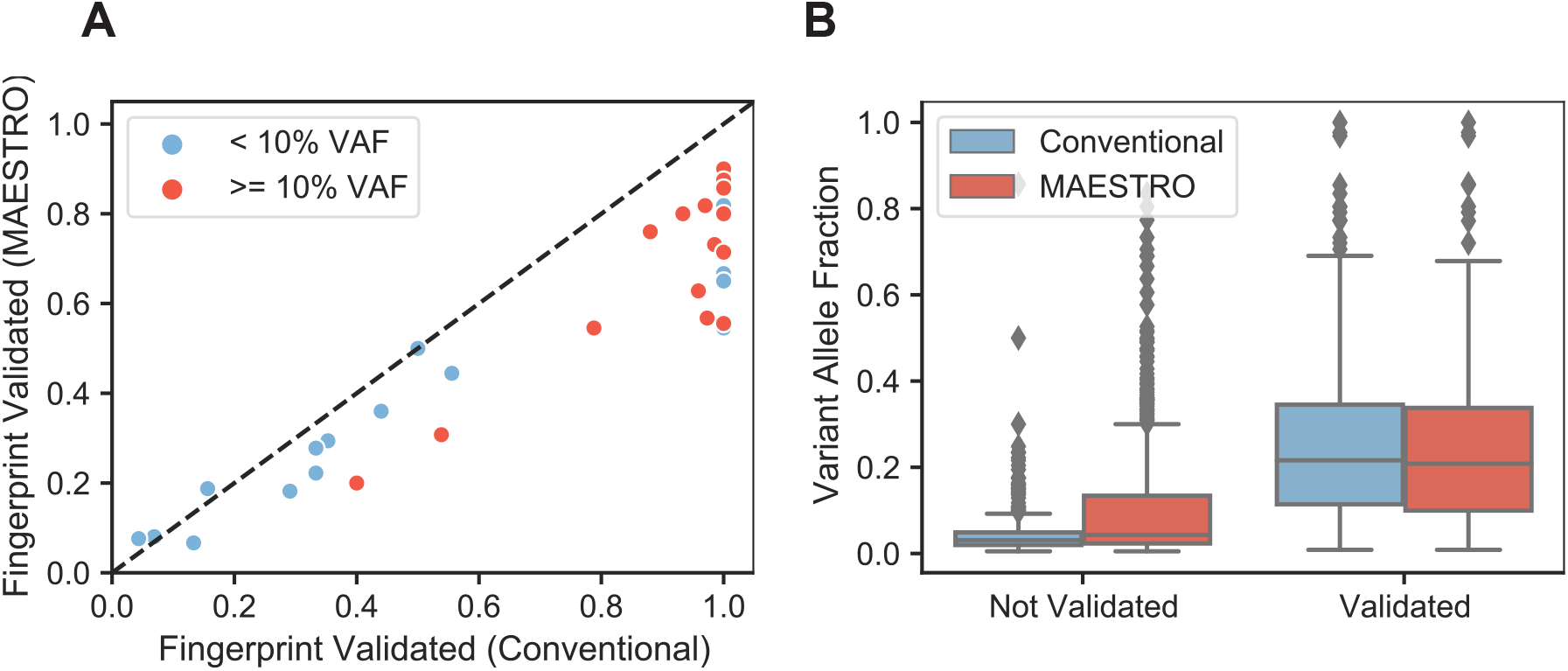
MAESTRO fingerprint validation of whole exome tumor samples. (A) Performance of 16x tumor fingerprints using both Conventional and MAESTRO. Mutations were called from the 16x tumor biopsies and both Conventional and MAESTRO fingerprints were created for all possible mutations from each tumor. The tumor biopsy libraries were captured with the Conventional and MAESTRO fingerprints and duplexes were sequenced. Fingerprints were split into two groups based on whether or not their original tumor VAF was < 10%. A mutation was considered validated if it was observed in the sequenced duplexes of the Conventional or MAESTRO sample. (B) Comparing variant allele fraction across all mutations from all Conventional and MAESTRO panels.

### MAESTRO could enable liquid biopsies to track up to 10,000 individualized mutations

To further characterize performance, and explore the feasibility to detect trace levels of ultra-rare mutations via liquid biopsy, we compared MAESTRO to conventional duplex sequencing for tracking 438 mutations in 18 x replicate 1/100k dilutions and 17 x replicate negative control samples. We used sheared genomic DNA from the same two cell lines described in the previous section to mimic cfDNA^8,34,38–42^ and isolated 20 ng for each replicate to reflect the cfDNA in typical 10 mL blood samples. These were intended to model the scenario for which (i) a limited mass of cfDNA fragments is drawn from the bloodstream, and (ii) at sufficiently low tumor fraction such that mutations are sparsely partitioned into each blood tube. At such ‘limiting dilution’, it becomes highly unlikely that the same mutation will be drawn in replicate samples and therefore, it is necessary to track many mutations^33,34^.

MAESTRO uncovered 81% (n=47/58) and 80% (n=4/5) of the mutant duplexes detected with Conventional across all 1/100k and negative control samples, respectively, using much less sequencing (**Supplementary Fig. 7A**). Most that were exclusive to Conventional in the 1/100k samples (n=6/11) were detected by MAESTRO but excluded by the noise filter. MAESTRO also uncovered an additional 52 and 16 mutant duplexes across all 1/100k and negative control samples, respectively, but most were near fragment ends, which proved less likely to be captured by Conventional in these experiments (**Supplementary Fig. 7B**). If we consider these differences, the concordance is nearly perfect (**Supplementary Fig. 7C**). However, for the rest of the study we do not remove the molecules that were less likely to be captured with Conventional. Importantly, MAESTRO detected significantly more mutations in the 1/100k samples than the negative controls (**Fig. 4A**, p=1.16E-5, Welch’s t-test). We also confirmed that without duplex error suppression, we would have been unable to resolve MRD at these limiting dilutions (**Supplementary Fig. 7D**).

**Figure 4:**
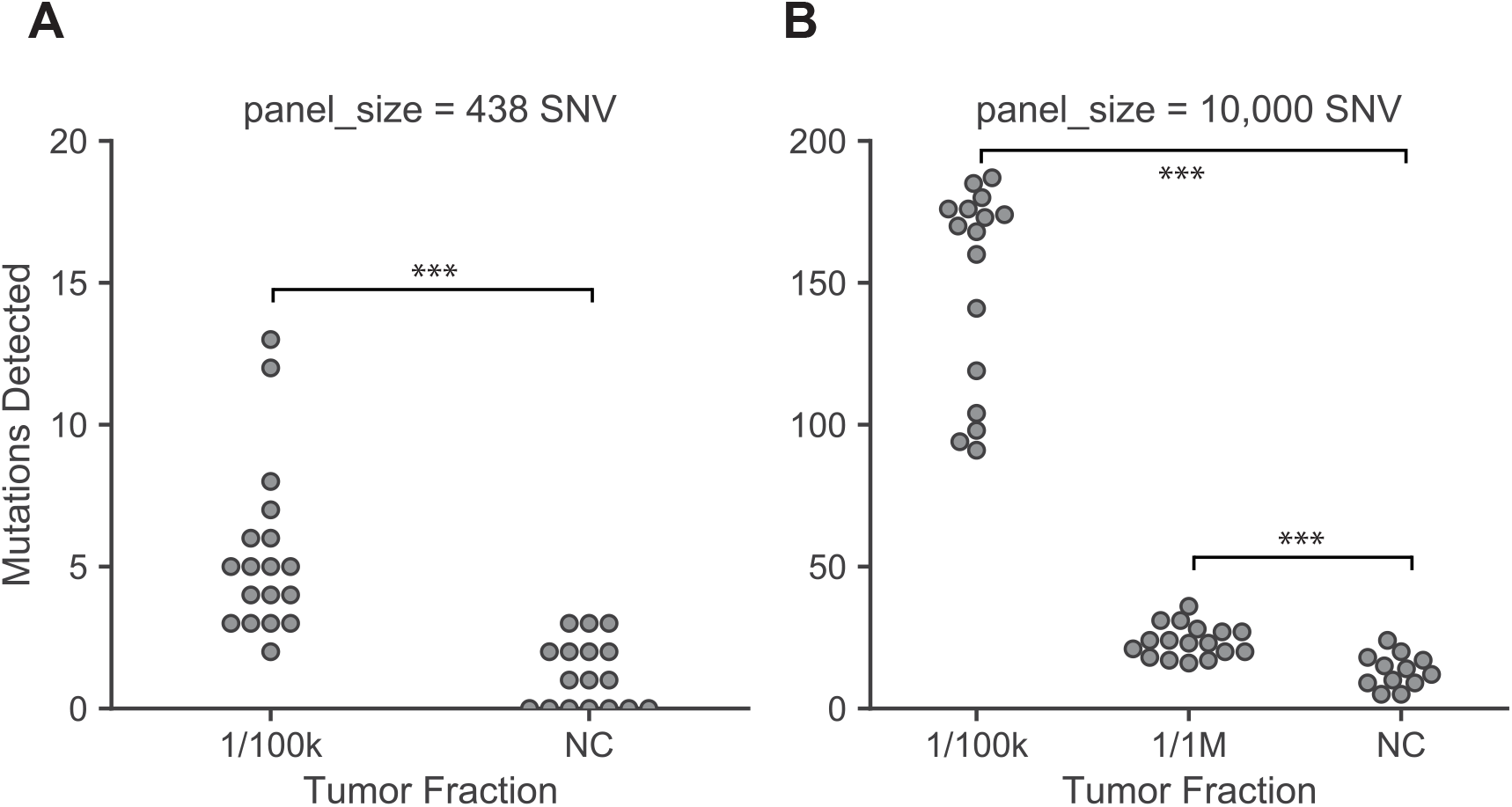
MAESTRO can detect signal above noise at 1/100k tumor fraction. (A) Mutations detected in MAESTRO using a 438 probe panel across 18 x biological replicates of a 1/100k dilution and 17 x biological replicates of a negative control. (B) Mutations detected in MAESTRO using a 10,000 probe panel across 16 x biological replicates of a 1/100k dilution, 17 x biological replicates of 1/1M, and 12 x negative controls. The Welch’s t-test was used to determine whether significantly more mutations were uncovered in each tumor dilution compared to the negative controls.

While MAESTRO provided comparable sensitivity and specificity using significantly less sequencing, the number of mutations detected at 1/100k dilution, of 438 tracked, was not much greater than the negative controls. We thus hypothesized that tracking even more mutations, e.g. 10,000—the typical number in a cancer genome^43^—could improve our signal-to-noise ratio and enhance MRD detection. Yet, this could only be done feasibly with MAESTRO, as Conventional would require >10 billion reads (~$20,000 on the Illumina HiSeqX) to saturate duplex recovery, in contrast to about ~100 million reads (~$200) with MAESTRO, in addition to other costs of sample preparation.

Applying MAESTRO to track 10,000 mutations in 16 x replicate 1/100k dilutions, 17 × 1/1M dilutions and 12 × negative controls, we observed a large increase in number of mutations detected in the 1/100k samples (median mutations=169, range 91 to 187), which was significantly higher than the negative controls (median 13 mutations, range 5 to 24, p=7.23E-11, **Fig. 4B**). Higher mutation counts were also observed in the 1/1M dilutions (median 23, range 16 to 36, p=7.47E-5), although further refinements are likely needed to enable reliable detection at 1/1M. Our results suggest that tracking thousands of genome-wide mutations provides a profound boost in our signal-to-noise ratio, which is likely to be crucial for tracking MRD and guiding treatment.

As for the mutations in the negative controls, we reason that these could either be (a) true mutations that arose spontaneously with each cell division, (b) cross-contamination when cell lines were cultured, or (c) technical artifacts that have yet to be overcome in duplex sequencing. While we cannot yet discern the source, the mutation counts were consistent with what was expected for scanning tens of millions of bases for potential mutation (10,000 mutations x few thousand haploid genomes of DNA) given the reported error rate of ~1×10^−6^ in duplex sequencing^13,34,37^. By retesting specific loci, we also verified that the majority would have been detected with conventional duplex sequencing (**Supplementary Fig. 8**), suggesting that most are not artifacts of the MAESTRO protocol.

### Tracking thousands of mutations from patients’ tumor genomes in cfDNA improves MRD detection

Considering the profound boost in our signal-to-noise ratio in dilution series, we sought to determine whether tracking all genome-wide tumor mutations could enhance MRD detection from cfDNA. For patients with some common, aggressive forms of breast cancer, standard care involves preoperative systemic chemotherapy for its utility in guiding subsequent response-based treatment^44,45^. We analyzed patients with breast cancer enrolled in a clinical trial (16 patients) of preoperative therapy **(Supplementary Fig. 9A)**, reasoning we could (i) determine the detectability of tumor-derived cfDNA at diagnosis, (ii) describe how cfDNA trends with clinical response over the course of treatment, and (iii) determine whether preoperative MRD testing could predict the presence of residual cancer in the surgical specimen.

Reasoning that genome-wide mutation tracking would be most useful in samples with low tumor fraction, we first tracked all exome-wide tumor mutations using a personalized cfDNA test built on conventional duplex sequencing^34^. We found that most patients had detectable circulating tumor DNA at diagnosis (median tumor fraction 0.00858, range 0 to 0.21, **Supplementary Fig. 9B**) and that a decrease in tumor fraction in cfDNA between the first two time points (T1, T2) trended with clinical response (**Supplementary Fig. 9C**) which is consistent with prior reports^27–29^. Yet, MRD was detected preoperatively (T4) using conventional duplex sequencing in only one of eight patients with residual disease at the time of surgery and in only one of five who experienced future distant recurrence. We chose the remaining four patients to explore whether genome-wide mutation tracking could enhance MRD detection.

For these four patients who had tested MRD-negative preoperatively but experienced future distant recurrence, we performed PCR-free whole-genome sequencing of their tumor biopsy specimens and blood normal DNA. We identified a median of 5575.5 (range 3385 to 8783) somatic mutations per patient and, using stringent criteria for probe design, we created one MAESTRO test comprising 55-58% of exonic mutations and 30-38% of intronic mutations from all patients (**Supplementary Fig. 10**). We applied the MAESTRO test to tumor and normal DNA and found 52% (range 41-56%) of probed mutations to be verified (**Supplementary Fig. 11**). We then applied the assay to all available cfDNA samples from all four patients, such that we assess all mutations in all patients, using the unmatched samples as controls for one another. By also applying MAESTRO tests to matched germline DNA from each patient, we further limited the potential impact of variants arising from clonal hematopoiesis.

We found that tracking all tumor mutations with MAESTRO uncovered more mutations per patient in cfDNA compared to Conventional (**Fig. 5A**) and detected no false mutations in any unmatched samples (**Fig. 5B**). Previous studies have shown that using > 1 mutation for MRD detection helps to protect against error^25,33,34^. We uncovered multiple tumor mutations preoperatively for two of the four patients, while observing profound signal enhancement in the earlier time points from all patients. These proof-of-principle results suggest that MAESTRO could enhance MRD detection by enabling all genome-wide tumor mutations to be accurately tracked in cfDNA.

**Figure 5:**
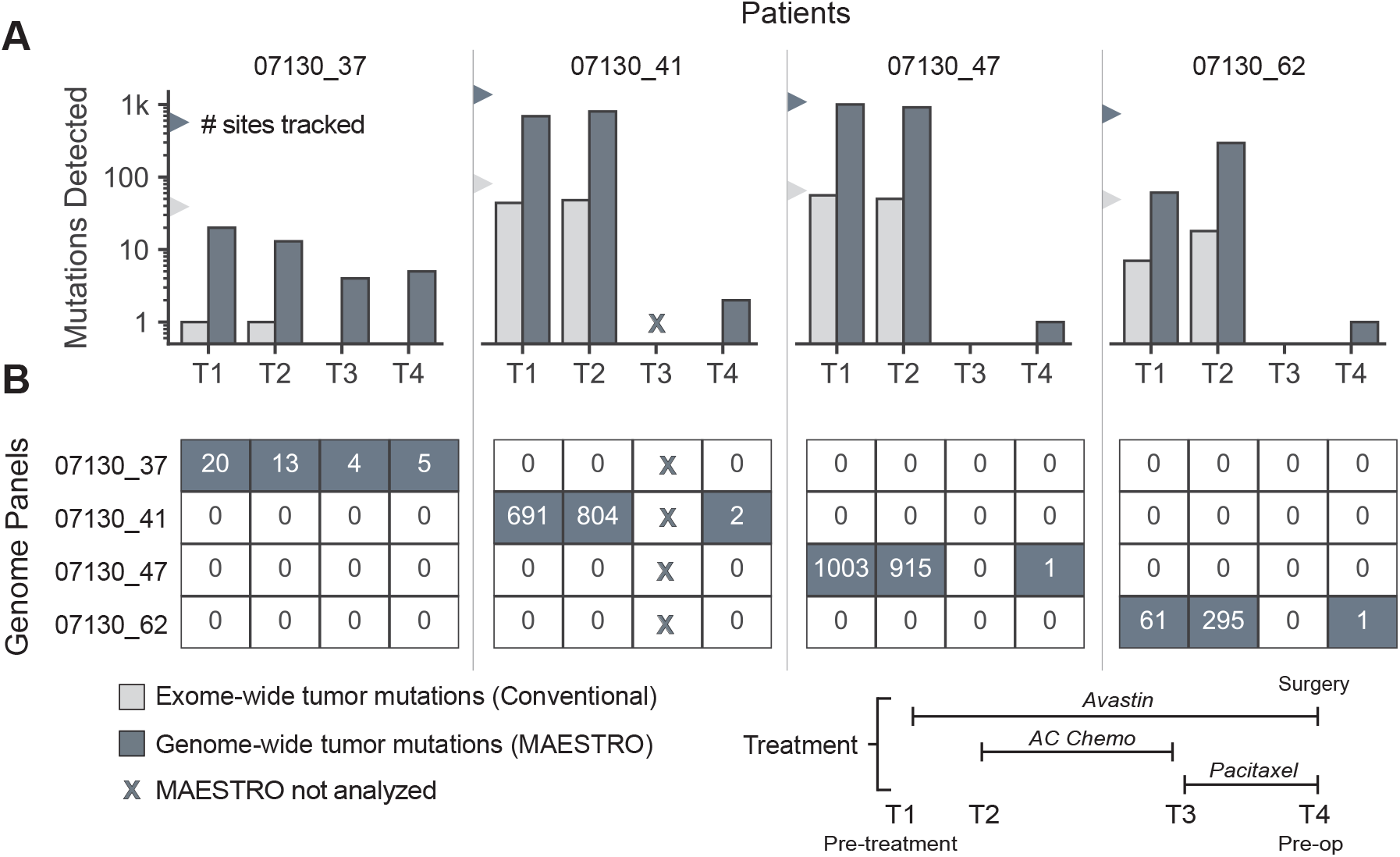
MAESTRO improves detection of MRD in pre-operative setting. (A) Genome-wide tumor mutations detected with MAESTRO compared to exome-wide tumor mutations detected with a personalized MRD test built on conventional duplex sequencing^34^. Fingerprint sizes for the two conditions are shown with triangles. Mutations from all patients were combined into a single panel for MAESTRO and the same panel was applied to all samples. (B) The heatmap shows mutation counts detected using MAESTRO with patient-specific mutations on the diagonal and highlights MAESTRO’s specificity.

## Discussion

In summary, we have demonstrated a simple and practical approach to extend the breadth, depth, accuracy, and efficiency of mutation tracking in clinical specimens. Our technique breaks the breadth-vs-depth ‘glass ceiling’ of DNA sequencing, enabling thousands of low-abundance mutations to be accurately tracked at low cost. This is likely to empower many types of biomedical research and diagnostic tests that demand accurate and efficient tracking of many rare mutations. For instance, we have shown that MAESTRO uniquely enables thousands of genome-wide tumor mutations to be tracked in liquid biopsies, and that this improves the detection of MRD after cancer treatment.

MAESTRO is the first method, to our knowledge, to simultaneously enrich and detect thousands of genome-wide mutations with high-accuracy sequencing. In a dilution series involving sheared genomic DNA, we demonstrated a median ~1000-fold enrichment from 0.1% VAF to nearly pure mutant DNA, which enabled us to detect most mutant duplexes using ~100-fold less sequencing. We showed that MAESTRO could track up to 10,000 distinct, low-abundance (< 0.1% VAF) mutations scattered throughout the genome. This is important because existing methods can scan for all possible mutations within consecutive bases (e.g. within the same amplicons or probed loci) but break down when it comes to tracking many mutations in non-overlapping regions, such as genome-wide tumor mutations. MAESTRO was designed to track predefined mutations—not for mutation scanning or discovery.

Our study is the first to track thousands of genome-wide tumor mutations from liquid biopsies, with sufficient breadth and depth to improve the detection of MRD. This is significant because (i) detecting MRD remains a significant unmet medical need, and (ii) while MRD detection correlates with the number of tumor mutations tracked in cfDNA^27,34,35^, existing techniques have had limited breadth or depth. For instance, cancer gene panels typically cover just a few mutations per patient^37^; patient-specific assays track tens to hundreds^27,33^; and whole-genome sequencing remains far too costly to apply beyond minimal depth^46^. Using MAESTRO, we found many more mutations detected at limiting dilutions such as 1/100k, from about 5 when 438 were tracked to almost 200 when 10,000 were tracked. Applying MAESTRO to patients undergoing neoadjuvant therapy for early-stage breast cancer, we detect significantly more when all genome-wide tumor mutations were tracked in comparison to all exome-wide mutations. With this improved sensitivity, we believe MAESTRO may also potentially benefit the postoperative and longitudinal detection of minimal residual disease. Bespoke genome-wide liquid biopsies reflect one potential application for MAESTRO. We have shown that tracking more mutations per patient improves the signal-to-noise ratio for MRD detection, suggesting that this could be valuable for the field. Yet, it remains to be determined whether this approach will outperform other existing tests, including epigenetic-based methods.

The profound signal enhancement that we observed for detecting MRD from liquid biopsies is likely to be important for guiding key treatment decisions such as to intensify therapy long before clinical recurrence, or to de-escalate treatment in a patient who does not have residual disease. For instance, our detection of hundreds of mutations at 1/100k limiting dilution could enable more confident determination of MRD status by placing less weight on any single mutation. This could help to overcome spurious mutations arising from clonal hematopoiesis. It could also empower new classification methods that leverage features such as fragment size that may be ‘less specific’ for any single mutation but informative when integrated across many mutations. While our approach requires whole-genome sequencing of each patient’s tumor and individualized probe design, the cost of each continues to decline, and biotinylation of oligonucleotides in-house can further help to limit costs (see **Methods**). We also expect that upfront costs could be amortized over many serial MRD tests, while being offset by large savings in sequencing required per test.

MAESTRO addresses a fundamental challenge in the mutation enrichment field by using molecular barcodes to discern true mutations from low-level errors that may also be enriched. Specifically, our DSC/SSC ratio filter is a novel advance that measures intrinsic noise within each sample, but two current limitations are (i) that it needs to be tuned, and (ii) that error-prone loci are discarded, which impacts sensitivity when these regions contain real mutations. One simple way to address this is to recapture MAESTRO-detected loci with probes that target both mutant and wild type, as we had done to confirm high specificity, but a better solution will be to recover all library molecules in the read family irrespective of mutant or wild type. We are currently working on a method that physically links top and bottom strands such that they can be recovered together.

Another limitation of mutation enrichment is that it may lose the ability to quantify mutation abundance. We are working to address this by incorporating internal controls to calibrate enrichment performance on a locus-by-locus basis, as well as probes against fixed sequences to estimate the total molecular diversity of the library and to confirm whether we have sequenced to saturation. We also focused on enrichment of point mutations, but expect that MAESTRO could also be useful for tracking other types of alterations such as insertions and deletions or structural variants. While tracking more mutations per patient could increase the number of unique cfDNA molecules sampled (and therefore, the detection limit for MRD)^27,35,37,46^, it will never be possible to detect MRD at tumor fractions below sequencing error rates. Accordingly, we opted to employ the most accurate sequencing method, duplex sequencing. In vitro and in silico methods exist to enrich circulating tumor DNA based upon size selection^47^ and preferred end coordinates^48^ but come nowhere near MAESTRO in terms of fold-enrichment.

In all, MAESTRO is a simple yet powerful approach to (i) convert low-abundance mutations into high-abundance mutations, and (ii) enable their detection with high-accuracy sequencing using significantly fewer reads. This means that it is no longer necessary to trade breadth for depth, or accuracy for efficiency, when tracking many low-abundance mutations in clinical samples. While we expect this to be useful in many ways, we are particularly excited about the ability to improve MRD detection, as this could lead to more precise care for millions of cancer patients.

## Methods

### Patients and Samples

All patients provided written informed consent to allow the collection of blood and/or tumor tissue and analysis of clinical and genetic data for research purposes. Patients with triple-negative breast cancer (TNBC) and a tumor size >1.5 cm were prospectively identified for enrollment into tissue analysis and banking cohorts (Dana-Farber Cancer Institute [DFCI] IRB-approved protocol 07130). Patients had plasma isolated from 20 cc blood in EDTA tubes and tissue sampling performed within six months of diagnosis. All patients completed the following course of neoadjuvant Phase II therapy: Bevacizumab x 1 dose; Doxorubicin/Cyclophosphamide x 4 cycles plus Bevacizumab; Paclitaxel x 4 cycles plus Bevacizumab. Blood draws were taken before each course. A Residual Cancer Burden (RCB) score was calculated after surgery. For those patients with sufficient tumor tissue, exome-sequencing identified mutations that we captured using our Conventional assay^34^. From within this cohort we identified four TNBC patients who had tested MRD-negative using the exome-wide panel but who experienced metastatic recurrence. For these patients, we applied MAESTRO to analyze genome-wide tumor mutations. HapMap DNA from NA12878 and NA19238 was purchased from Corielle. This research was conducted in accordance with the provisions of the Declaration of Helsinki and the U.S. Common Rule.

### Defining Mutations to Track

For the HapMap panels, VCF files were taken from the Genome in a Bottle Consortium^49^ (NA12878) and 1000 Genomes project^50^ (NA19238). Sites specific to NA12878 were subsampled to create MAF files and were subsequently run through probe design to create the 438 and 10,000 SNV (single nucleotide variant) fingerprints.

Tumor DNA was extracted from fresh-frozen tumor samples. All patients’ tumor DNA underwent whole-exome sequencing to identify trackable mutations for conventional capture. Of the four patients selected for MAESTRO, tumor DNA underwent PCR-free whole-genome sequencing. Illumina output from whole-genome sequencing was processed by the Broad Picard pipeline and aligned to hg19 using BWA. We ran the GATK best practices workflow on the Terra platform to detect somatic SNVs and indels in our deep whole-genome sequencing data using tumor/normal calling (<u>see Terra workflow</u>). We subset the somatic mutation calls to only SNVs and passed the candidate SNVs for tracking to our probe design pipeline. By sequencing each patient’s tumor and normal to adequate depth we can avoid tracking variants arising from clonal hematopoiesis.

### Probe Design

Mutations in MAF (mutation annotation format) were first checked for specificity in the reference genome to filter out potential mapping artifacts. The resulting filtered MAF was then used as input into probe design. Conventional probe design was performed on the filtered MAF as previously described^34^. For MAESTRO probe design, along with the mutation file, initial probe length (default = 30 bp), annealing temperature (default = 50°C), and Δ*G* range (default = −18 to −14 kcal/mol) were used as input. For Δ*G* and melting temperature calculations, we use the annealing temperature, [Na^+^] = 50 mM, [Mg^2+^] = 0 mM, and [DNA] = 250 nM. An initial sequence was designed for the given length with the mutation at its center. If the sequence was within the specified Δ*G* range, it proceeded through the subsequent design steps, otherwise the sequence length was adjusted until it fell within the range. A modified BLAST was performed where the melting temperature for each hit was calculated and if it was less than the annealing temperature, it was removed. If there were 10 or greater pass-filter BLAST hits, we attempted to redesign the sequence using a sliding window. This resulted in the mutation being offset from the center of the sequence, but still provided good enrichment. The sequence with the minimum BLAST hits was then chosen. All sequences were output in a tab-delimited file, and the results were filtered based on length, GC content, Δ*G*, and the number of BLAST hits before ending up with the final panel design (**Supplementary Fig. 1**).

### In-house Biotinylation of Probe Panel

Patient-specific oligo pools ordered from Twist Bioscience contained universal forward and reverse primer binding sites. Amplification of the oligo pool was performed using an internally biotin-modified forward primer containing a dU base directly 5’ to the biotinylated dT and an unmodified reverse primer containing a BciVI recognition sequence at its 3’ end. The PCR product was purified using Zymo’s DNA Clean & Concentrator-25 columns. Two micrograms of biotinylated, double-stranded product were sequentially subject to the following 100 μL one-tube enzymatic reaction: 40 units BciVI for 60 minutes at 37°C; 10 units Lambda Exonuclease for 30 minutes at 37°C followed by 20 minutes at 80°C; 7 units USER Enzyme for 30 minutes at 37°C (NEB)^51^. Zymo’s Oligo Clean & Concentrator columns were used to purify short, single-stranded, biotinylated probes for hybrid capture.

### DNA Extraction and Library Construction

Healthy gDNA from two HapMap cell lines, NA12878 and NA19238, were sheared to 150 bp fragments using a Covaris E220/LE220 Ultrasonicator. Sheared DNA was quantified using the Quant-iT Picogreen dsDNA assay kit on a Hamilton STAR-line liquid handler. Tumor fraction dilutions were created by spiking sheared gDNA from NA12878 (“tumor”) into NA19238 (“normal”) at 0, 1:1K, 1:10K, 1:100K, and 1:1M tumor fractions. All libraries were constructed with 20 ng sheared gDNA using the Kapa Hyper Prep Kit with custom dual-index duplex UMI adapters (IDT). These UMI adapters allowed tracking of the top and bottom strand of each unique starting molecule despite rounds of amplification. Processing of patient blood samples followed the same protocol as previously described^52^. Germline DNA (gDNA) was extracted from either buffy coat or whole blood using the QIAsymphony DSP DNA Mini kit and sheared. Cell-free DNA (cfDNA) was extracted from plasma using the QIAsymphony DSP Circulating DNA Kit. cfDNA and gDNA libraries were constructed in the same manner as HapMap DNA.

In cases where there was insufficient library remaining for a subsequent capture, 200 ng of library was subject to additional rounds of PCR to generate workable mass (>1 μg) for hybrid capture using KAPA’s library amplification primer mix. In cases where technical replicates of the same library were needed, libraries were reindexed using a new set of P5/P7 indices (IDT).

### MAESTRO capture

Hybrid capture using biotinylated, short probe panels was performed using xGen Hybridization and Wash Kit with xGen Universal Blockers (IDT) using a protocol adapted from Schmitt, et al^36^. Each hybrid capture contained 1 μg of library and 0.75 pmol/μL of MAESTRO probes (IDT or Twist Bioscience), using wells in the middle of the 96-well plate to prevent temperature fluctuations. The hybridization program began at 95°C for 30 seconds. This was followed by a stepwise decrease in temperature from 65°C to 50°C, dropping 1°C every 48 minutes. Finally, the plate was held at 50°C for at least four hours, making the total time in hybridization 16 hours. Heated wash buffer was kept at 50°C (lid temp 55°C) and heated wash steps were performed at 50°C. After the first round of hybrid capture, 16 cycles of PCR were applied. The product was subject to a second round of hybrid capture using half volumes of Cot-1 DNA, xGen Universal Blockers, and probes. This was followed by another 16 cycles of PCR. Apart from these differences, MAESTRO double capture was performed using the same protocol as outlined in Parsons, et al^34^. Final captured product was quantified and pooled for sequencing on an Illumina HiSeq 2500 (101 bp paired-end reads) or a HiSeqX (151 bp paired-end reads) with a target raw depth of 10,000 x per site.

### Conventional Capture

The following described protocol was outlined previously in Parsons, et al^34^. Hybrid capture using a panel consisting of patient-specific (i.e. germline informed), biotinylated 120 nt probes was performed using the xGen Hybridization and Wash Kit with xGen Universal Blockers (IDT). For each Conventional capture reaction, libraries were pooled up to 6-plex with 500 ng input each and 0.56 to 0.75 pmol/μL of probe panel was applied (IDT). The hybridization program began at 95°C for 30 seconds. This was followed by 65°C for 16 hours. Heated wash buffer was kept at 65°C (lid temp 70°C) and heated wash steps were performed at 65°C. After the first round of hybrid capture, 16 cycles of PCR were applied. The product was subject to a second round of hybrid capture using half volumes of Cot-1 DNA, xGen Universal Blockers, and probes. This was followed by another 8 cycles of PCR. Final captured product was quantified and pooled for sequencing on an Illumina HiSeq 2500 (101 bp paired-end reads) or a HiSeqX (151 bp paired-end reads) with a target raw depth of 1,000,000 x per site.

### Quantification of Library Conversion Efficiency by ddPCR

To quantify library conversion efficiency, a ddPCR assay was designed to target the flanking adapter regions. Only fragments with successful double ligation were exponentially amplified within the QX200 ddPCR EvaGreen Supermix (Bio-Rad). Varying DNA inputs into LC (3ng, 10ng, 20ng, 50ng) were tested for their varying conversion efficiencies and adjusted to an unligated control. The results are shown in **Supplementary Table 1.**

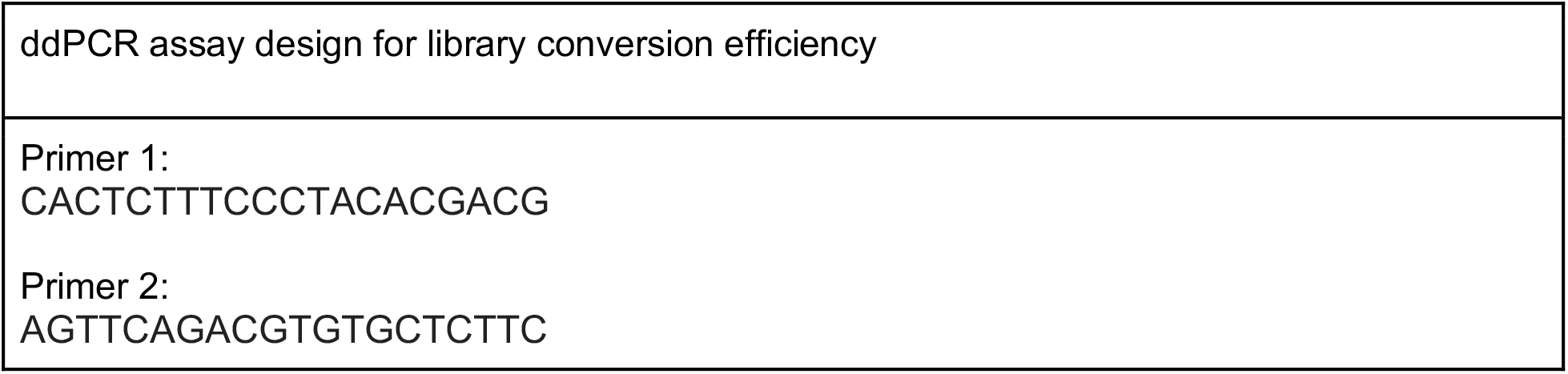

### Quantification of Probe Capture Efficiency by ddPCR

To quantify probe capture efficiency, a ddPCR assay was designed to target a homozygous mutation site chosen from the 438 SNV HapMap fingerprint (see ddPCR assay design below). Conventional and MAESTRO hybrid capture was performed on pure tumor Hapmap gDNA libraries, with all waste streams collected from washes. The total number of mutant molecules into hybrid capture and lost during hybrid capture were quantified by using the designed ddPCR assay^33,34^. Probe capture efficiencies were determined using the equation below. The results are shown in **Supplementary Table 2**.

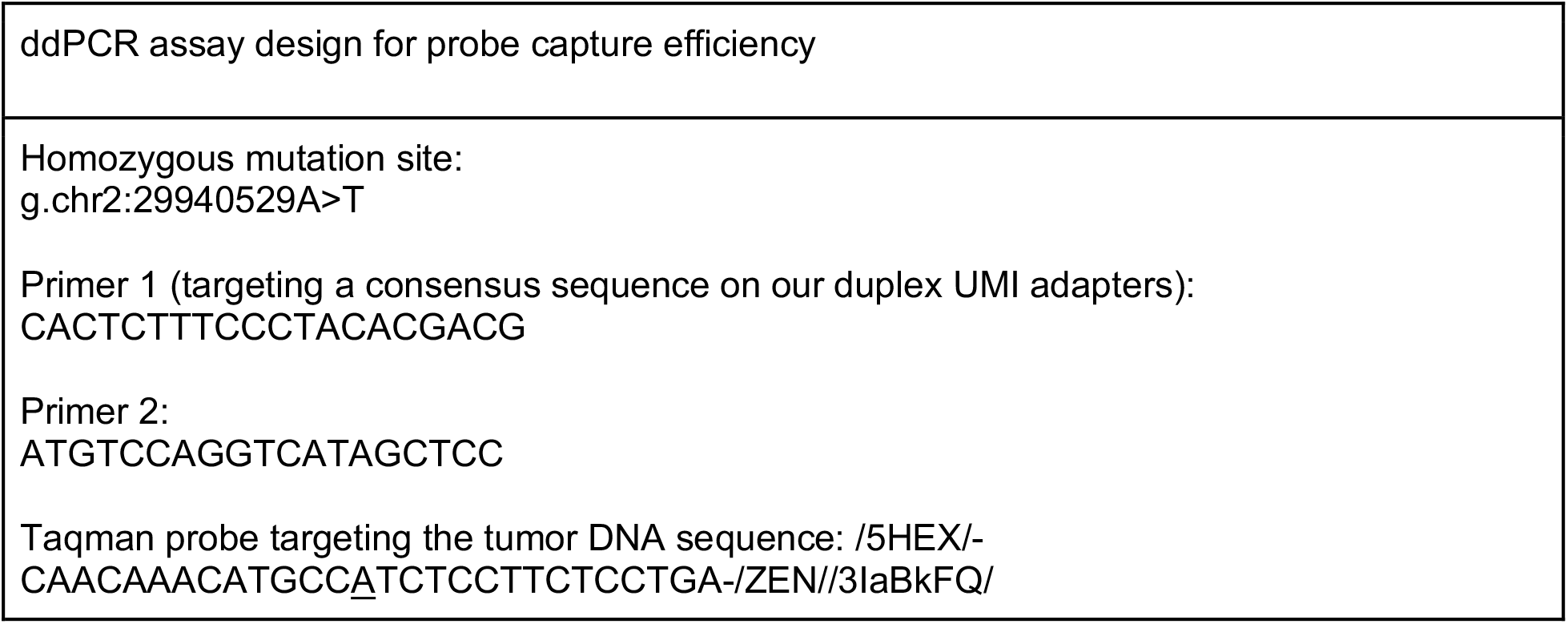

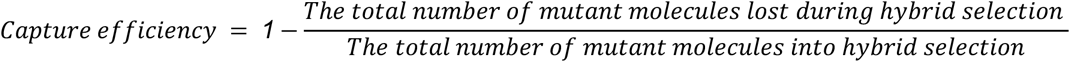

### Sequencing and Data Analysis

Sequencing and pre-processing of BAM files followed a similar protocol as previously described^33,34^ with the following changes. Before grouping reads by UMI, read groups were added to samples from the same library and samples were merged into a single BAM. This ensured identical molecules found in different samples were given the same family ID from Fgbio’s GroupReadsByUmi (Fulcrum Genomics). The resulting BAM was then pushed through GroupReadsByUmi and split afterwards by the added read group tag. The split BAMs were then passed through the consensus calling workflow. Consensus BAM files were indel realigned using GATK 4 before calling mutations using custom scripts. Noise filtering based on DSC/SSC ratio (total mutant DSCs / total mutant SSCs) was performed on all MAESTRO samples. For mutation calling in clinical samples, we used both the matched tumor and normal. We required that each mutation be seen in the tumor and not in the normal in order for the mutation to be considered. Processing of BAM files was automated using a Snakemake^53^ workflow (**Supplementary Fig. 12**).

### Miredas Minimal Residual Disease Analysis Scripts

A suite of scripts (Miredas) was used for calling mutations and creating metrics files. In the Snakemake workflow, MiredasCollectErrorMetric uses the duplex BAM file to describe the number of errors and calculates errors per base sequenced. MiredasDetectFingerprint uses the duplex BAM file to call mutations and MiredasDetectFingerprintSsc uses the single-stranded BAM file to call mutations. This single-stranded output of MiredasDetectFingerprintSsc is used along with the duplex MiredasDetectFingerprint output to create DSC/SSC ratios.

### VAF/Recall

Raw VAF was calculated using the single strand consensus BAMs as consensus bases are more reliable compared to raw sequenced bases and help correct for PCR bias. We opted to use the single strand consensus BAMs rather than the duplex BAMs as we wanted to retain the majority of sequenced reads - with duplex sequencing, more than 50% of reads can be lost due to support only being observed on one strand. For each site, a pileup was created from the single strand consensus BAM and read bases were compared to the called bases in the MAF file. Each base was categorized as reference (REF), alternate (ALT), or OTHER and the consensus family size (number of reads contributing to the consensus) was added to the site’s read counts. Raw VAF could then be calculated by comparing the number of ALT reads to the total reads (REF + ALT + OTHER) for each site. This raw VAF measurement is important for determining the efficiency of sequencing the ALT base, but may not be an accurate readout of true variant allele fraction due to PCR bias. To address this, we have included duplex VAF in **Supplementary Table 4**, where duplex VAF is calculated using the consensus duplex fragments rather than family size as used in raw VAF.

To assess recall, the duplex consensus BAM files were used. Our consensus calling workflow gives source molecules the same family ID, so two samples from the same library have many overlapping molecules. Recall was calculated by looking at the overlap of duplex families between two samples (oftentimes a Conventional sample and a MAESTRO sample). See **Supplementary Fig. 3B** for an example.

### Noise Filter

Four replicate negative controls were created from the same source library via reindexing as described in **DNA Extraction and Library Construction**. The replicates were captured using the 10,000 SNV MAESTRO panel. For each targeted site with ALT molecules present in any of the replicates, a DSC/SSC ratio was calculated by summing all ALT supporting duplexes and dividing by the total ALT supporting single strand consensus molecules. Targets with ALT duplexes present in more than one replicate were considered “shared” whereas targets with ALT duplexes present in a single replicate were marked as “exclusive”. A single DSC/SSC ratio was chosen that maximized the number of targets shared while minimizing the number of exclusive targets.

### Probe Spike-in Experiment

MAESTRO capture was performed with a 10,000 SNV panel applied to negative control HapMap samples. Prior to post-capture PCR, ten MAESTRO probes selected randomly from the 10,000 SNV panel and synthesized by IDT were added at 1000x concentration. This created a worst-case scenario to test the hypothesis that excess probe can create new mutant molecules by extending from real molecules, specifically during post-capture PCR (see **Supplementary Fig. 5A** for a schematic of this hypothesis). The usual post-PCR cleanup removed all excess probes. Second capture proceeded in the same manner.

### Tumor Fraction Estimation

Methods for calculating tumor fraction were previously described^33,34^ but some changes were made for use with MAESTRO. In a conventional sample, the full wildtype and mutant diversity is available and can inform tumor fraction. This is important as our tumor fraction methods currently rely on first calculating allele fraction (ALT depth / total depth) for all sites. In MAESTRO samples, we often have full mutant diversity, but wildtype molecules have been depleted. Because enrichment is not perfect, for each panel we have some targets that retain the full diversity of wildtype. We currently leverage this imperfect enrichment to estimate what the total potential depth of the sample is (how many cells likely contributed to the cfDNA library). This estimated depth is applied to all targets which allows us to calculate allele fraction (without considering copy number alterations) and subsequently tumor fraction. **Supplementary Fig. 13** shows this strategy and how it compares to actual tumor fractions. We are aware that these methods are not perfect in their current state, but believe that advances in quality control (ie testing for a handful of germline SNPs to measure unique duplexes per loci) could further improve tumor fraction estimation from enriched samples.

## Data Availability

All sequencing data will be deposited into a controlled access database such as dbGaP.

## Code Availability

Software tools will be made available via GitHub.

## Acknowledgments

First and foremost the authors would like to acknowledge the patients and their families for their contributions to this study. The authors would also like to thank the generous support from the Gerstner Family Foundation. This project was supported in part by the Bridge Project, a partnership between the Koch Institute for Integrative Cancer Research at MIT and the Dana-Farber/Harvard Cancer Center (DF/HCC). The authors also acknowledge support from National Institutes of Health grants R33 CA217652 and R01 CA22187

