## Supplemental Figures for "MAESTRO affords ‘breadth and depth’ for mutation testing"

### MAESTRO affords ‘breadth and depth’ for mutation tracking in clinical specimens

### Supplementary Figures

January 21, 2021

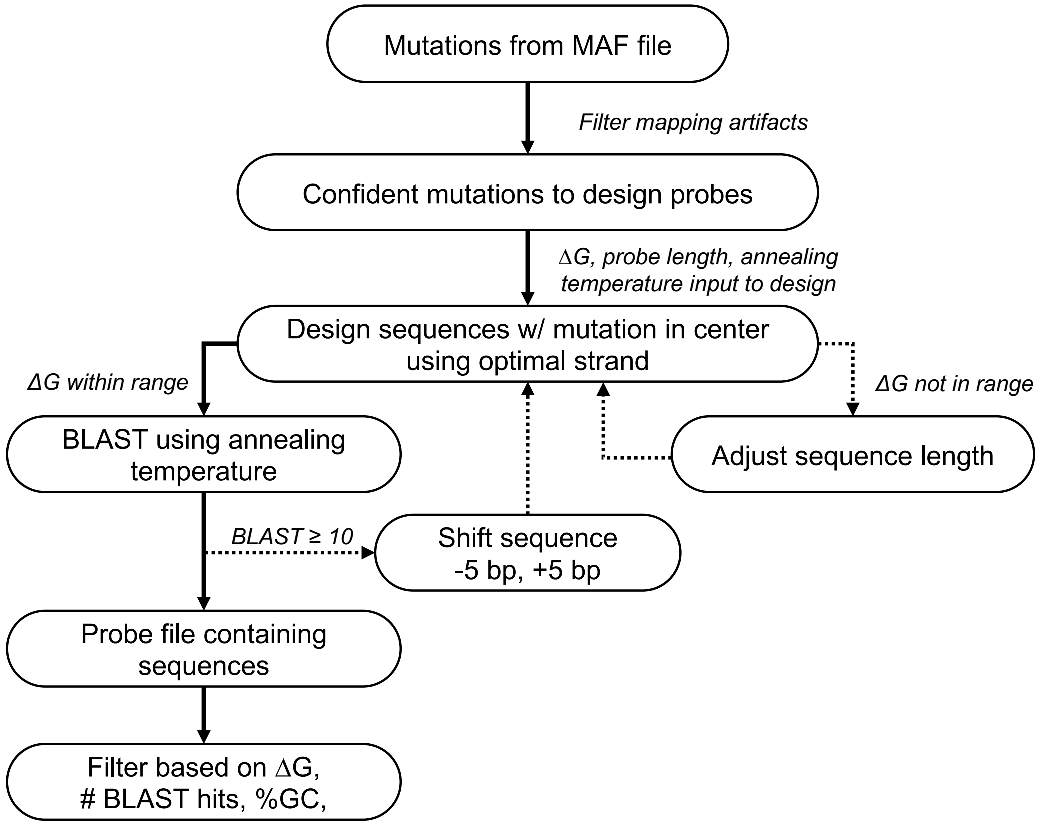

**Supplementary Figure 1**: **Probe design overview.** See Probe Design section of Methods.

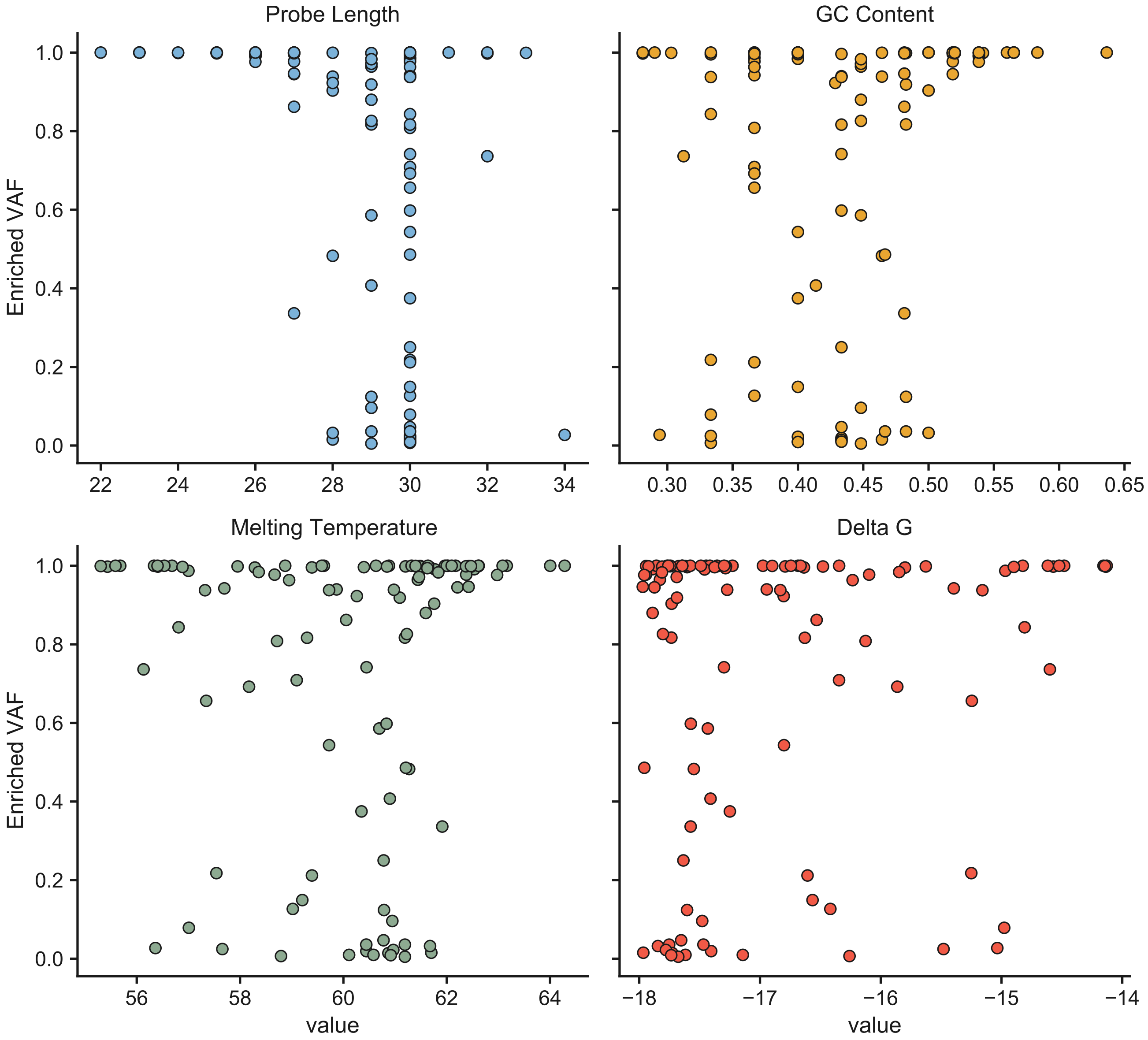

**Supplementary Figure 2**: **Probe characteristics effect on enrichment.** Showing results from the 1/1k dilution samples where each data point is a probe within the capture panel. Enriched VAF is plotted as a function of different probe sequence characteristics.

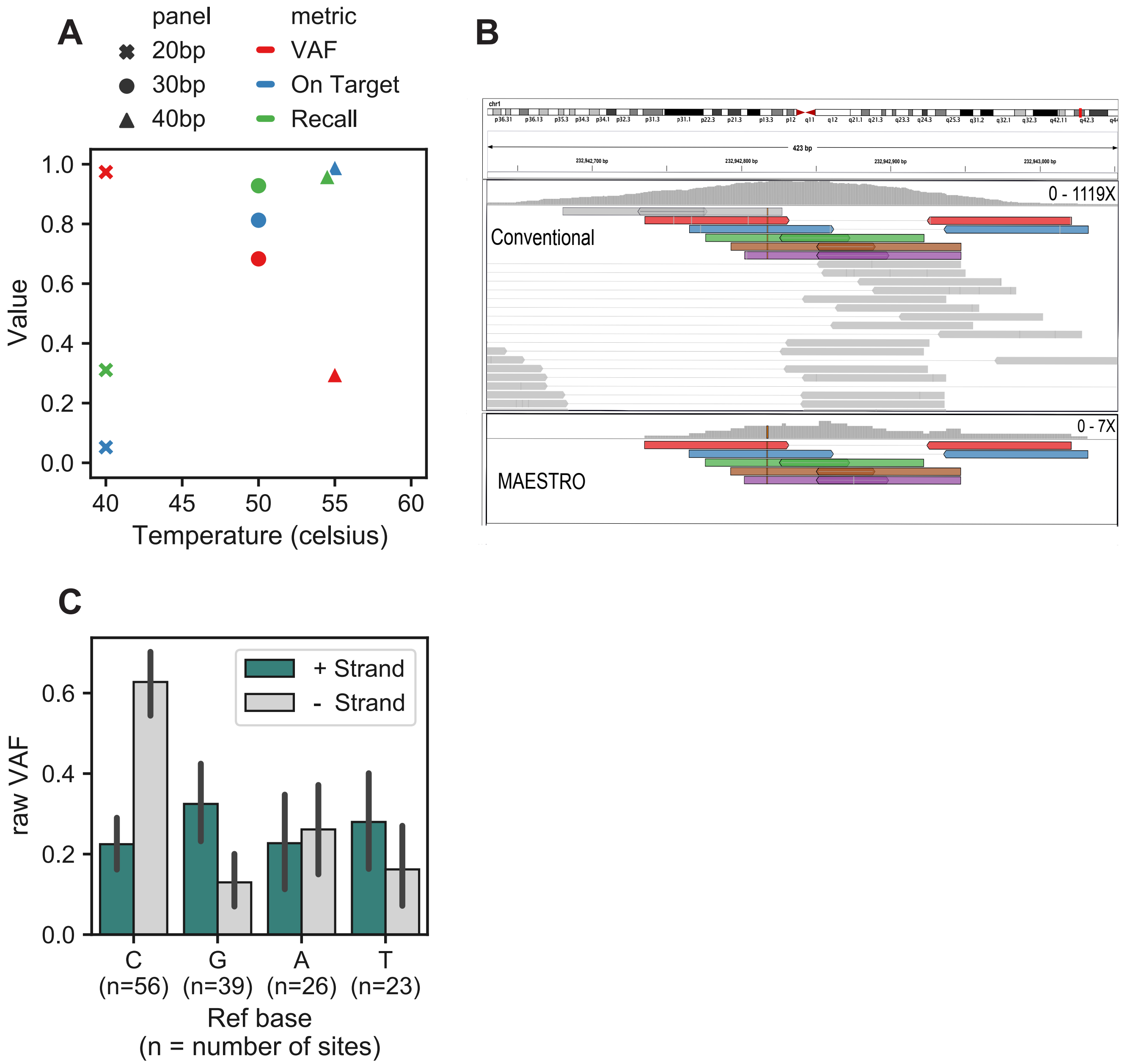

**Supplementary Figure 3**: **Probe and hybridization optimization.** (A) Effect of varying probe length and hybridization temperature on enrichment performance measured using variant allele fraction (VAF), on target fraction, and recall. All temperatures were tested for each probe length, but only the best performing temperature is shown. Data points for VAF and recall show mean across 20 sites whereas on target is calculated once per sample (total bases on target / total bases sequenced). (B) IGV screenshot showing an example of recall. Here, the same sample was captured using conventional and MAESTRO and identical source duplexes are colored. Recall in this example is 5/6 as 5 of the conventional duplexes were seen in the MAESTRO condition. (C) When designing probes, either the top or the bottom strand can be used. There will be different mismatches between the probe and wildtype base depending on which strand is chosen. Here, for each reference base across 144 sites, a MAESTRO probe was designed for either the top or the bottom strand and VRF performance is shown. When the reference base is a "C" it is beneficial to design for the negative strand. In all other cases, the positive strand is optimal. Showing mean with error bars representing 95% confidence interval. Interestingly, we did not observe equal and opposite magnitude raw VAF changes when swapping strands of C and G reference base probes. We believe this may be due to differences in probe characteristics (i.e. delta G, length) for each base category but further investigation is needed.

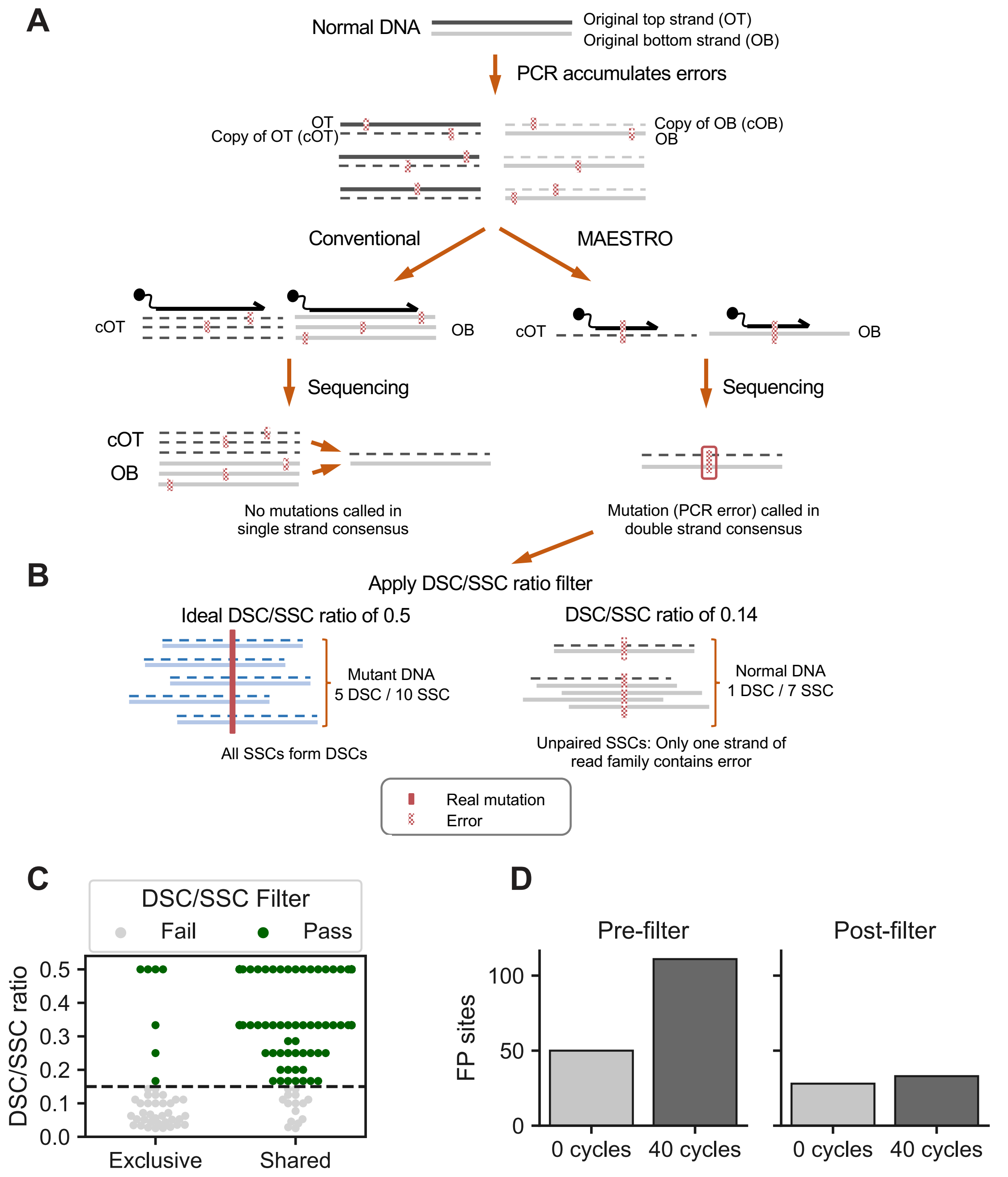

**Supplementary Figure 4**: **Tunable MAESTRO filter to correct for PCR errors.** (A) Library molecules accumulate polymerase errors during PCR. In conventional capture, PCR errors are suppressed by sequencing through all molecules at a given site, mutated or not. Errors can be corrected because they are seen spuriously and do not pass single strand consensus (SSC). With MAESTRO probes, PCR errors at the target base are also captured and sequenced. If an unmutated library molecule acquires the same PCR error on fragments derived from both the top and bottom strand of the same starting molecule, a false mutation is called even after double strand consensus (DSC). (B) In order to filter rare PCR errors that make it through duplex consensus, we can apply a DSC/SSC filter. To verify a mutation is real, most SSCs at the mutant site must be involved in forming a DSC (ideal DSC/SSC ratio of 0.5). Because PCR errors are impartial to read family, an accumulation of unpaired SSCs without accompanying DSC support signals a false mutation. (C) MAESTRO locus specific noise filter applied to four replicate negative controls. Molecules shared in at least two replicates are shown as well as molecules exclusive to one replicate. After applying the noise filter the majority of exclusive molecules are removed and shared molecules are retained. (D) Comparison of a sample with no added cycles of PCR to the same sample but with 40 added cycles before and after incorporating the DSC/SSC noise filter. Samples in both C and D used the 10,000 SNV panel.

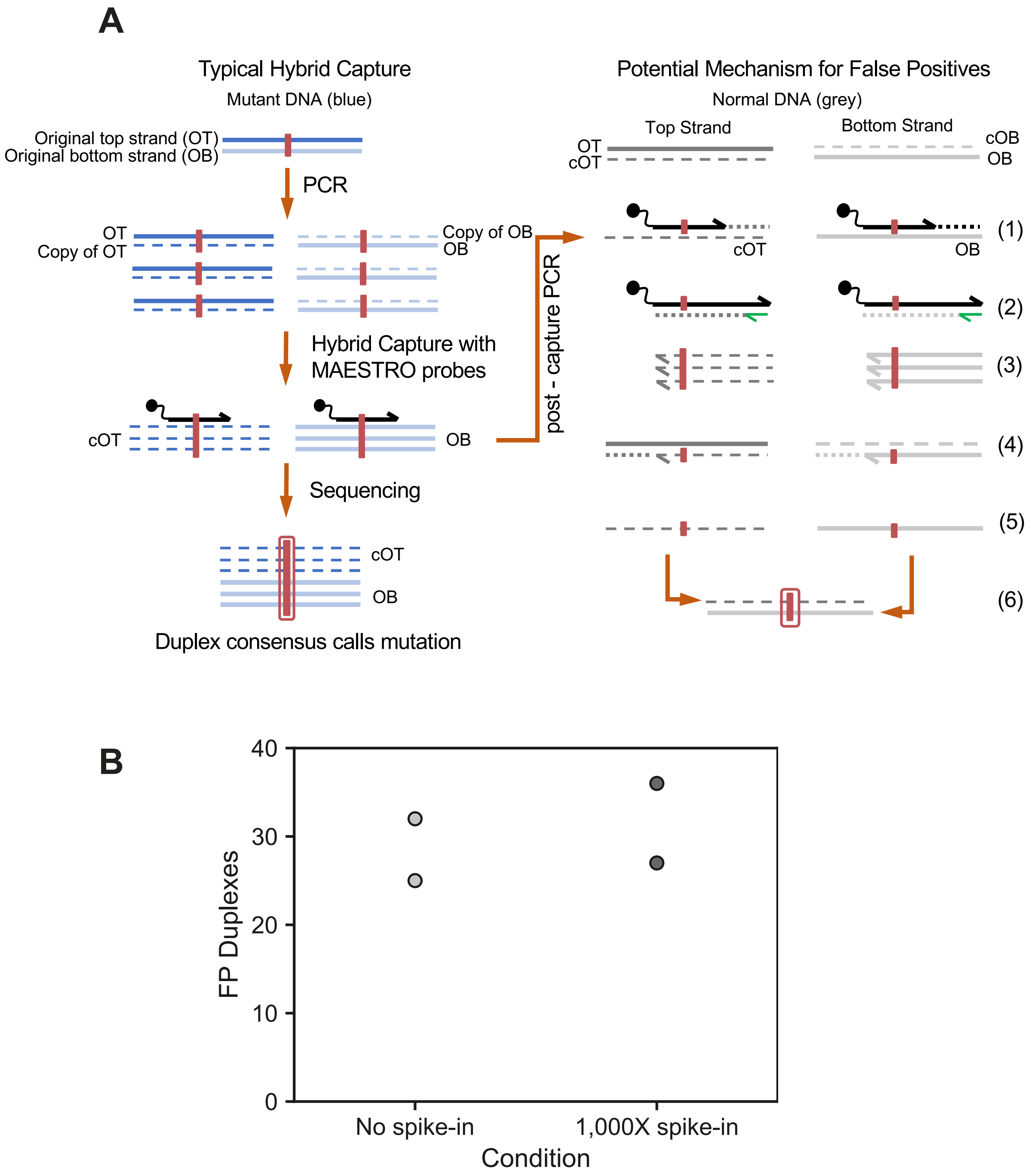

**Supplementary Figure 5**: **Probe spike-in experiment.** (A) Schematic showing how probes contain mutation of interest and may have the ability to create mutant duplexes. In order for a mutation to be called after duplex consensus, evidence must be seen in molecules derived from both the original top and bottom strand. During the 16 cycles of PCR performed after capture, a MAESTRO probe could bind to a non-mutant fragment and extend (1). This extended probe could be amplified in the next few rounds of PCR using the Illumina primers present in post-capture PCR (2). The copied products contain the mutation but are not able to be sequenced (3). These products can then bind to another unmutated fragment and extend (4). This creates a mutant molecule with both adapters intact that can be sequenced (5). This can result in a false-positive during duplex consensus if the same events happen on the other strand (6). (B) Capture was performed using the 10,000 SNV MAESTRO panel on two replicate negative control samples (no spike-in) and compared to the same negative controls with 1,000X the standard concentration of ten MAESTRO probes added prior to both post-capture PCRs (1,000X spike-in).

A

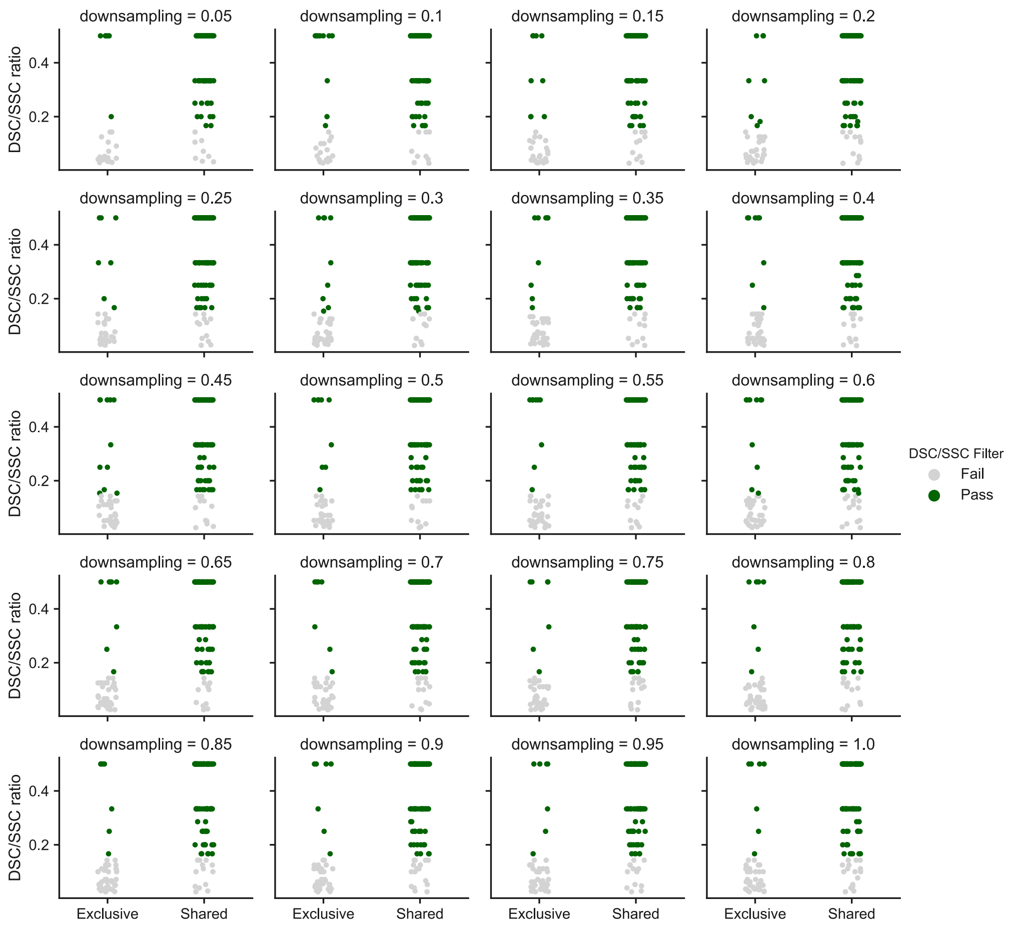

B

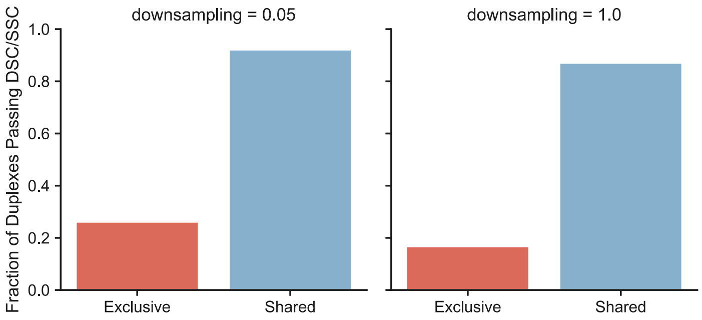

**Supplementary Figure 6**: **Downsampling DSC/SSC ratio.** (A) MAESTRO locus specific noise filter applied to four replicate negative controls with downsampling ranging from 1.0 (full sequencing depth used) down to 0.05 of the original depth. The samples and definitions are as described in Supplementary Fig. 4C. (B) A direct comparison of the fraction of duplexes passing DSC/SSC ratio filter at 1.0 (full sequencing depth) compared to 0.05 of the original depth.

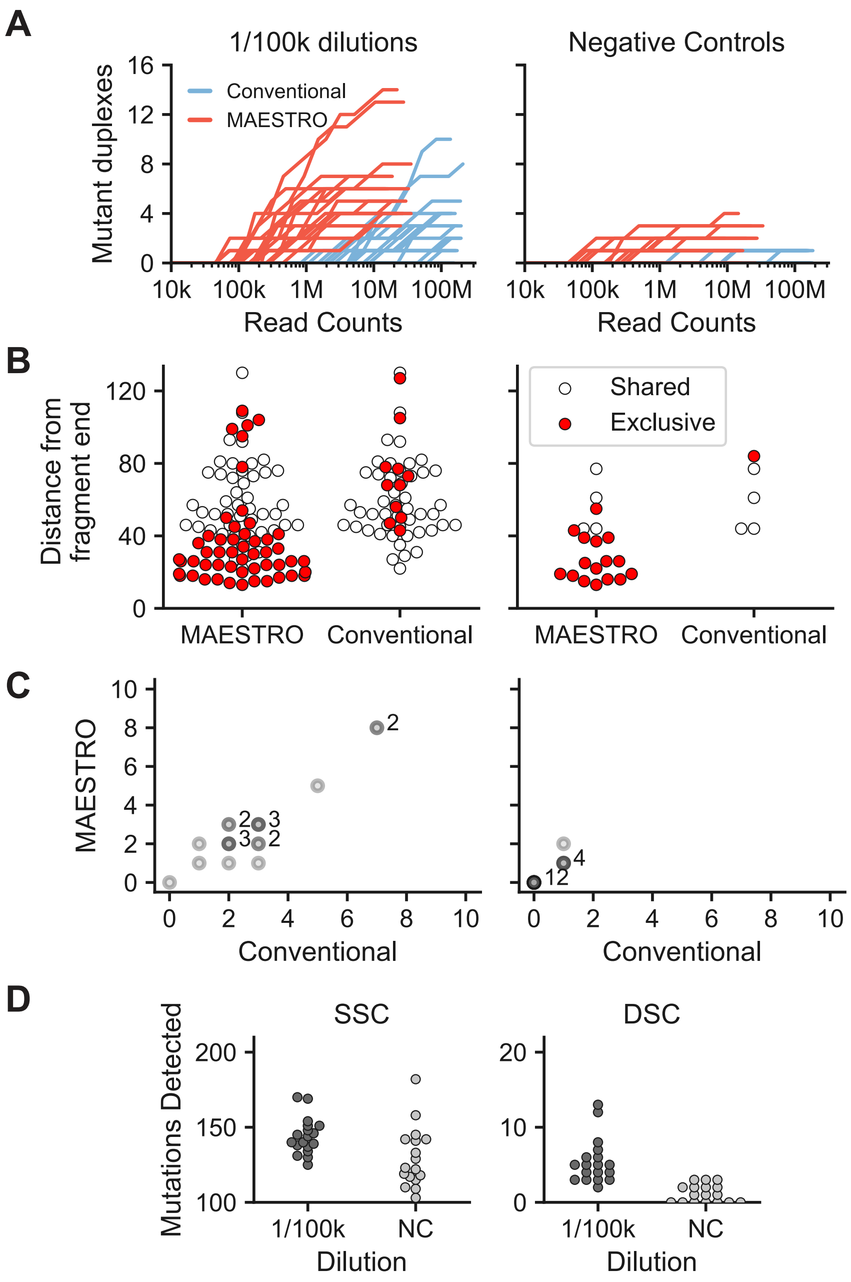

**Supplementary Figure 7**: **Benchmarking 1/100k dilutions.** Figures A, B, C, and D all use 18 x replicates of a 1/100k dilution and 17 x replicates of a negative control with a 438 SNV panel. (A) Comparison of downsampling curves resulting from applying conventional duplex sequencing and MAESTRO to the same replicate samples. (B) Distance from mutation site to fragment end (using the end closest to the mutation) shown for all mutant molecules uncovered with conventional and MAESTRO. Molecules with mutation near fragment ends were efficiently captured with MAESTRO probes but were not captured with conventional probes. (C) Removing molecules near fragment ends compensates for the different capture efficiencies of conventional and MAESTRO probes and results in high concordance between the two methods. Each axis contains the mutation counts seen across replicates. Points are shaded based on the number of replicates that overlap and any datapoint with more than one replicate is annotated with a number. (D) With single strand consensus sequencing, many additional mutations are uncovered in the negative control making it difficult to distinguish signal from noise.

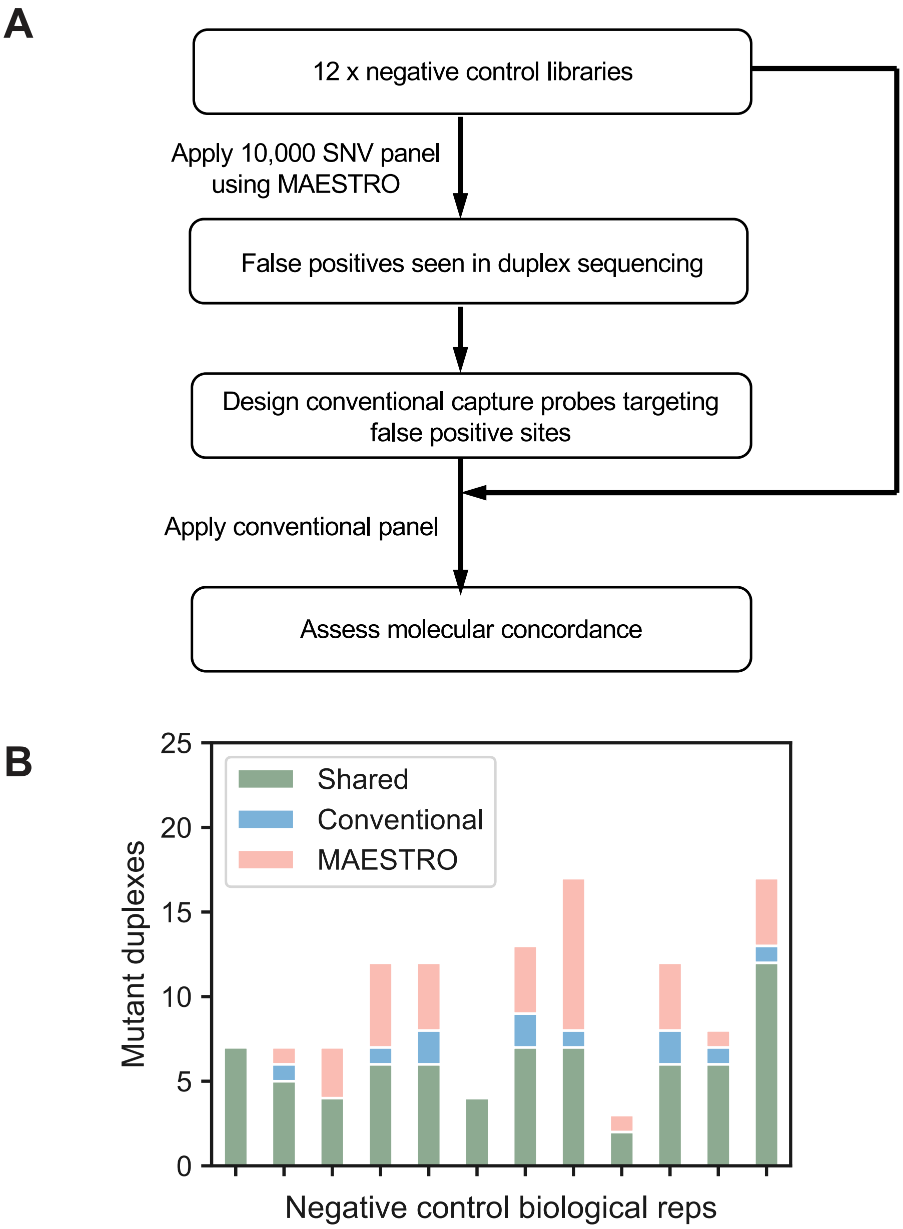

**Supplementary Figure 8**: **Validation of false positives in negative controls.** (A) Validation experiment design. (B) Duplex molecular concordance of false positives seen across 12 negative controls with conventional duplex sequencing and MAESTRO.

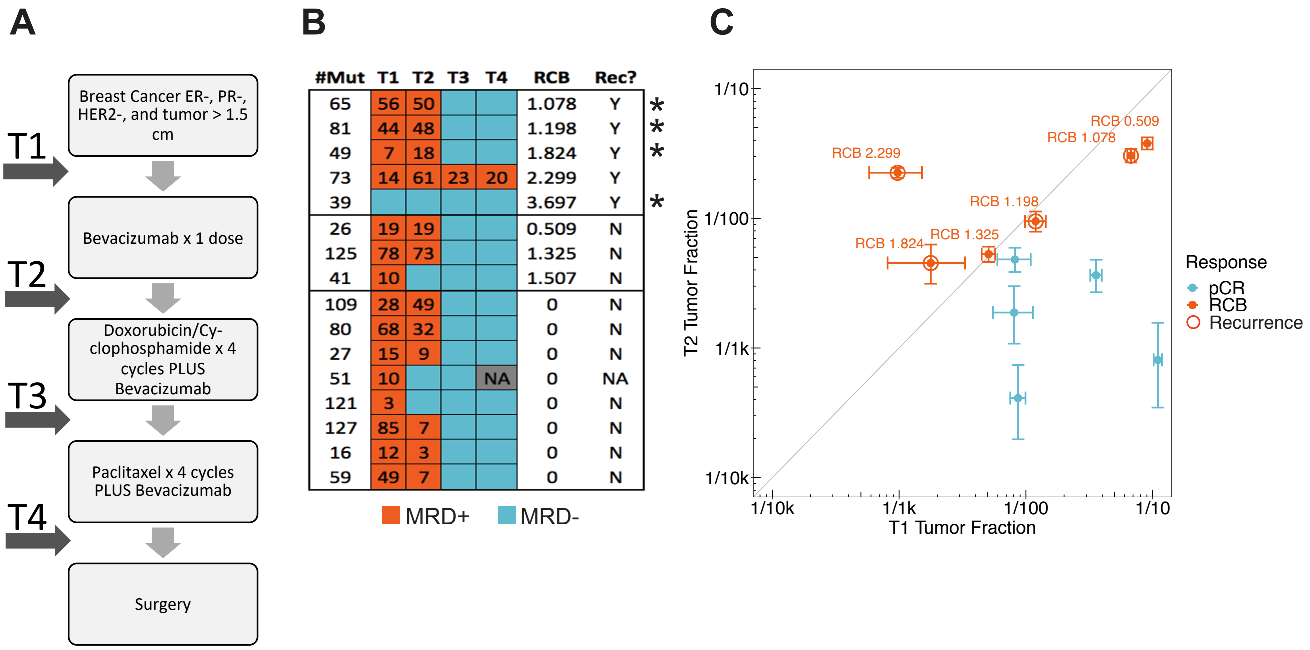

**Supplementary Figure 9**: **MRD testing in a Phase II study of preoperative doxorubicin and cyclophosphamide followed by paclitaxel with avastin in triple-negative breast cancer.** (A) Treatment course for patients from diagnosis to surgery with time of blood draw annotated. (B) Whole-exome sequencing of patients' tumor biopsies was performed, and individualized MRD tests were applied using conventional duplex sequencing to serial cfDNA time points as previously described (Parsons et al). MRD status (>=2 mutations detected) is indicated. Stars denote the four patients selected for more extensive testing with MAESTRO, results of which are shown in Fig 4A. (C) Comparison of tumor fractions from T1 and T2 blood draws. Data points are colored by pathological complete response or patients having residual cancer burden. Circles indicate patients that experienced recurrence. Error bars indicate 95% confidence intervals.

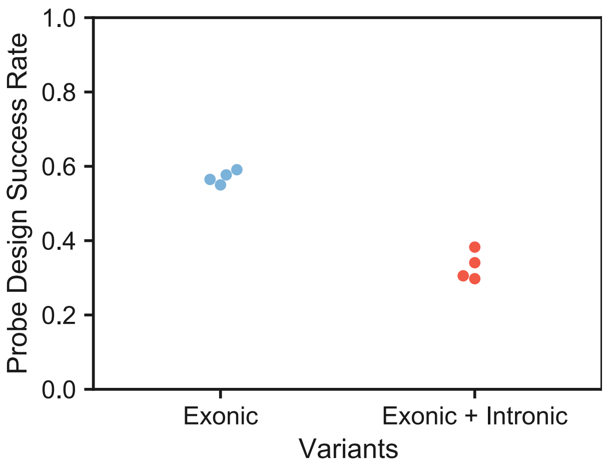

**Supplementary Figure 10**: **Probe design success rates.** Probe design success rate for the 4 patient-specific fingerprints analyzed in Fig. 5. Here, "Exonic" mutations were derived from whole exome sequencing of the tumor whereas "Exonic + Intronic" were from the combined output of whole exome and whole genome sequencing of the patient's tumor.

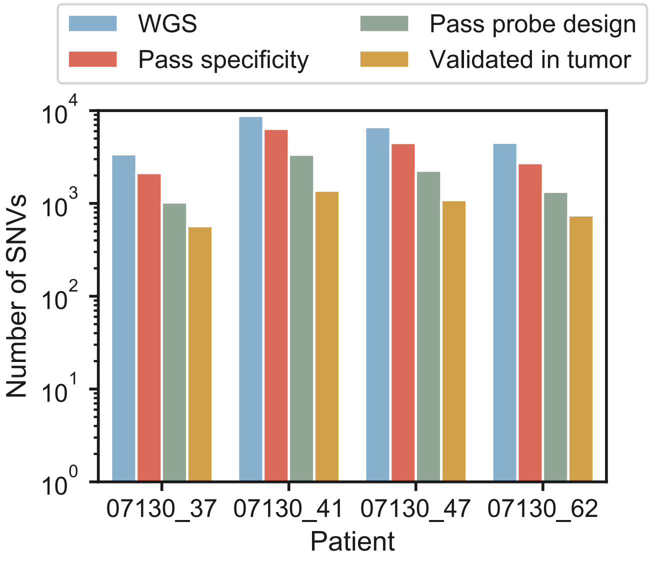

**Supplementary Figure 11**: **Somatic SNV counts and validation using patient's tumor DNA.** The total SNV counts from WGS is shown for each patient along with the total number of SNVs that pass our specificity filter that ensures good mappability. Next is the total number of SNVs that pass MAESTRO probe design and lastly are the total counts of mutations that were validated in each patient's tumor DNA.

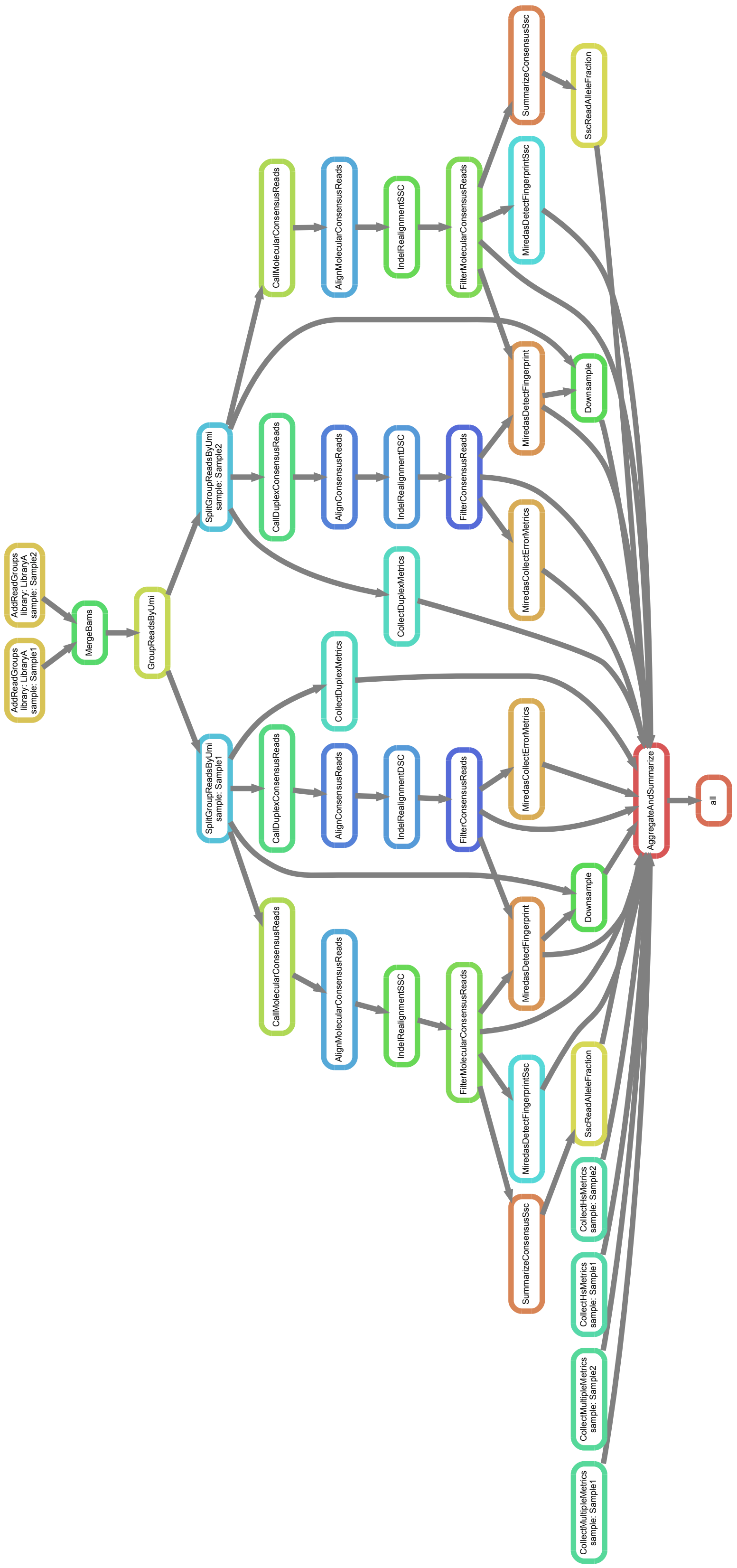

**Supplementary Figure 12**: **Analysis workflow DAG from Snakemake.** In the workflow shown, "library" is the original source library before capture. "Sample" is the library following capture (could be MAESTRO or Conventional). Samples from the same original source library are merged first so that common molecules are given the same unique family identifier.

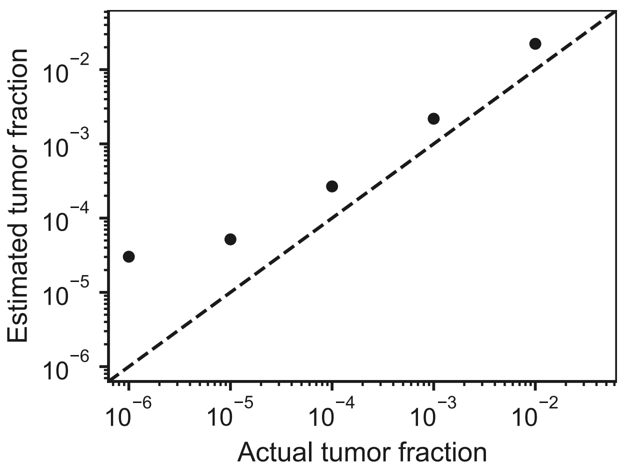

**Supplementary Figure 13**: **MAESTRO tumor fraction estimation.** Estimated tumor fraction compared to the actual tumor fraction for a spike-in tumor dilution series (see Methods, DNA Extraction and Library Construction). Estimated tumor fractions were calulated as described in the Tumor Fraction Estimation section of the methods.
